# Creating resistance to the whitefly *Bemisia tabaci* in cassava through RNAi-mediated targeting of multiple insect metabolic processes

**DOI:** 10.64898/2026.02.23.707345

**Authors:** Narayanan Narayanan, Rekha A. R. Swamy, Jackson Gehan, Tira Jones, Shelly Lazar, Dor Wintraube, Esther Yakir, Oren Hasson, Adam Lampert, John Colvin, Nigel J. Taylor, Shai Morin, Osnat Malka

**Author notes:** Narayanan Narayanan and Rekha A. R. Swamy equally contributed to this work. Correspondence: Nigel J. Taylor, Shai Morin, Osnat Malka.

## Abstract

It is commonplace in East Africa for 100% of cassava fields to be infected with Cassava mosaic disease (CMD) and/or Cassava brown streak disease (CBSD), resulting in annual losses of more than US$1.25 billion and reduced food and economic security for farming households. The vector of both diseases is the African cassava species of the whitefly *Bemisia tabaci*. Since the late 1990s, there has been an unprecedented increase in whitefly populations, to the extent that they are referred to as “super-abundant”. Research efforts since the late 1990s has focused mainly on developing plant resistance to the viral pathogens and paid scant attention to understanding the root causes of disease epidemics or the control of whitefly infestation. Here, we aimed at developing long-term whitefly-control solutions using an *in-planta* RNA interference (RNAi) approach. First, transcriptome analysis identified candidate genes that play ‘key’ roles in whitefly biology: osmoregulation, sugar metabolism and transport, symbiosis with endosymbiotic bacteria and detoxification of phytotoxins. Then, fifteen RNAi inverted repeat constructs were produced, designed to target the candidate genes and 140 independent transgenic lines were generated in cassava variety NASE 13. Whole plant bioassays showed insecticidal activity of transgenic plants, reaching 58% lethality for adults within 7 days and 75-90% lethality of nymphs after 25 days, compared to control plants. Target genes were confirmed to be downregulated by up to 2.5-fold in adult whiteflies and nymphs. We used population dynamics modelling to predict the potential of the RNAi technology to control whiteflies under field conditions in East Africa.

## Introduction

Cassava is recognized as a strategic food-security crop for smallholder farmers, particularly in sub-Saharan Africa where it supplies over half of caloric intake for about 200 million people and it is cultivated predominantly on farms smaller than one hectare (Obong’o *et al*., 2024). The importance of cassava for food security stems from its high carbohydrate yield, tolerance to drought and poor soils, and capacity to remain in the ground for extended periods, which together make it an “insurance crop” that buffers households against climatic and market shocks (Adebayo, 2023). The contribution of cassava to small-holder food security is increasingly constrained by viral diseases, vectored by the cassava-whitefly *Bemisia tabaci*. Since the 1990s, an unprecedented increase in the abundance of the cassava-whitefly, sub-Saharan Africa 1 (SSA1) and SSA2 species (Mugerwa *et al*., 2021a), has occurred across the cassava growing regions of East and Central Africa (Colvin *et al*., 2004; Legg *et al*., 2014; Macfadyen *et al*., 2018; Macfadyen *et al*., 2021). The cassava whitefly is responsible for vectoring the causal viruses of Cassava mosaic disease (CMD) and Cassava brown streak disease (CBSD). CBSD and CMD are the most important diseases of cassava in Africa, causing estimated production losses as high as 47%, equivalent to US$1.25 billion annually (Mugerwa *et al*., 2021b). In addition to vectoring virus diseases, super-abundant whitefly populations can cause up to 40% yield loss through direct feeding, and from reduced photosynthesis caused by black moulds (sooty mould) that develop on leaves as a result of whitefly-produced honeydew (Omongo *et al*., 2012).

To date, efforts to combat the effects of cassava whitefly have focused mostly on developing cassava varieties with robust resistance to the viral pathogens, with significantly less attention given to host plant resistance against the whitefly itself (Haggar *et al*., 2020; Ndunguru *et al*., 2009). Both must be achieved if the detrimental impact of this pest is to be negated. Breeding programs have had some success in selecting new varieties with elevated tolerance to whiteflies (Atim *et al*., 2024, Sam *et al*., 2024). However, these varieties do not combine elevated tolerance to whiteflies with other agronomic traits, such as high storage root yields, good dry matter content and the organoleptic qualities important for widespread adoption by farmers (Manze *et al*., 2021). It is also known that whitefly populations evolve to overcome tolerance (Tadmor *et al*., 2022). New and robust sources of resistance to cassava whiteflies are therefore required for breeders to pyramid with disease resistance in farmer-preferred germplasm.

We report here a biotechnological approach utilizing plant-expressed RNA interference (*in-planta* RNAi) to target genes involved in multiple processes essential to the cassava-whitefly life cycle. Like other hemipteran insects, whiteflies can ingest dsRNAs from phloem tissues, which trigger RNAi responses in the insect’s gut and downregulate the target genes (Rosa *et al*., 2018; Zhu and palli, 2020). Three previous studies have demonstrated the potential of using transgenic, plant-mediated RNAi to control whiteflies. Thakur *et al*. (2014) observed an enhanced level of *B. tabaci* resistance in transgenic *Nicotiana tabacum* plants expressing dsRNA targeting the vacuolar type H^+^ ATPase (*v ATPase A*) gene. Transcripts levels of *v ATPase A* in *B. tabaci* feeding on plants expressing this dsRNA was reduced by 62%, leading to 84% adult mortality after six days. Eakteiman *et al*. (2018) targeted the glutathione S-transferase (GST) detoxification gene, *BtGSTs5*. *B. tabaci* feeding on *Arabidopsis thaliana* plants expressing dsRNA derived from *BtGSTs5* showed significant down regulation of this target gene in their gut. An associated delay in nymph development by up to 3.5 days was observed, leading to significant reduction in population build-up. In the third case, Xia *et al*. (2021) targeted a glucoside malonyltransferase gene (*BtPMaT1*) which enables *B. tabaci* to neutralize phenolic glucosides. Adults feeding on transgenic tomato plants significantly reduced *BtPMaT1* expression and resulted in almost 100% adult mortality.

Previous genomic and transcriptomics analyses by our group and others (Jing *et al*., 2016; Luan *et al*., 2015; Luo *et al*., 2017; Malka *et al*., 2018; Malka *et al*., 2021) enabled us to develop four strategies for insecticidal RNAi activity. These target essential processes and pathways in the cassava-whitefly: osmoregulation, carbohydrate homeostasis, symbiosis and detoxification. First, to impair osmoregulation (ability to tolerate high osmotic pressure of ingested phloem-sap), we targeted an aquaporin (*AQP*) that mediates water cycling across the gut (Shakesby *et al*., 2009), and three GH13 α-glucosidases (*SUCs*) that contribute to osmotic balance via sucrose isomerization/oligomerization (Jing *et al*., 2016; Luo *et al*., 2017; Mathew *et al*., 2011; Wintraube *et al*., 2025). Second, to downregulate carbohydrate homeostasis and thereby reduce available energy and stress tolerance, we silenced genes encoding UTP-glucose-1-phosphate uridylyltransferase (*UGP*) and trehalose-6-phosphate synthase (*TPS*), which are central to UDP-glucose, trehalose, and glycogen biosynthesis (Thompson, 2003), as well as a sugar transporter (*ST*) required for sugar flux across membranes. Third, because *B. tabaci* relies on the obligate endosymbiont ‘*Candidatus* Portiera aleyrodidarum’ for the synthesis of essential nutrients, we targeted five horizontally acquired insect genes expressed in the bacteriocytes: arginosuccinate lyase (*argH*), chorismate mutase (*CM*), diaminopimelate decarboxylase (*LysA*), diaminopimelate epimerase (*DapF*), and biotin synthase (*BioB*), involved in the synthesis of arginine, phenylalanine, lysine, and biotin (Kaweesi *et al*., 2024). Finally, to compromise detoxification of plant defence phytotoxins, we silenced three cassava-whitefly genes coding for detoxification proteins: two UDP-glucosyltransferases (*UDPGT*s) involved in phase II conjugation and a gene coding for an ATP-Binding Cassette (ABC) transporter that exports neutralized compounds (Despres *et al*., 2007).

Inverted repeat expression cassettes were generated from sequences targeting the above genes and their expression was directed to phloem tissues in transgenic cassava plants using a phloem-specific promoter. Whole plant bioassays enabled assessment of insecticidal impact on adults and nymphs. Data generated was incorporated into population dynamics modelling to assess potential of these RNAi strategies to suppress build-up of cassava whitefly populations in the field under different regional cropping mixtures of RNAi-protected and non-protected varieties.

## Materials and Methods

### dsRNA design

dsRNA sequences were designed using the Snapdragon platform (Hu *et al*., 2021; https://www.flyrnai.org/cgi-bin/RNAi_find_primers.pl). The dsRNA regions were selected using an siRNA-design workflow in which high-quality candidate 21-mers were first predicted (DSIR), mapped back to the target transcript, and then used to place each dsRNA within regions of higher predicted siRNA density, with preference (when feasible) for regions proximal to the start codon. To screen for inter-species off-targets, all possible overlapping 21-nt k-mers were generated, using a 1-nt sliding window (splitter -size 21 -overlap 20), for each designed dsRNA sequence. Non-target organism datasets (RefSeq-based datasets) were searched with USEARCH against both strands, counting only exact matches (0 mismatches; -maxdiffs 0). A panel of 10 reference species was selected to represent key relevant non-target groups in East African cassava agroecosystems, including humans, domestic animals, pollinators, and the host plant cassava. Specifically, the following RefSeq assemblies were used: *Homo sapiens* (GCF_000001405.40), *Bos taurus* (GCF_002263795.3), *Sus scrofa* (GCF_000003025.6), *Canis lupus familiaris* (GCF_011100685.1), *Ovis aries* (GCF_016772045.2), *Capra hircus* (GCF_001704415.2), *Gallus gallus* (GCF_016699485.2), *Manihot esculenta* (GCF_001659605.2), *Apis mellifera* (GCF_003254395.2) and *Frieseomelitta varia* (GCF_011392965.1). Across all comparisons, no 21-bp perfect matches were detected in any of reference species, indicating the absence of major off-target candidates. All designed dsRNA sequences are available in Table S1.

### Production of RNAi constructs for plant transformation

RNAi gene constructs DWF10-17 and DWF55–68 (Table S2) were generated in which the *AtSUC2* promoter drives expression of inverted repeats of sense and antisense sequences separated by the *pdk* intron (Figure S1A) (Rosche *et al*., 1994). The binary vector p8384, a modified derivate of p5000 (Chauhan *et al*., 2015), was used as the backbone. Both the cassette and vector were digested with *SbfI* and *AscI* (NEB), ligated, and used for cloning. T-DNA cassettes, containing an *AtSUC2* promoter, inverted repeat and NOS terminator, were synthesized and cloned into the p5000 vector. The gene cassettes were excised with *XhoI/SbfI* and the *AtSUC2* promoter with *XhoI/AscI*, p5000 linearized with *SbfI/AscI* and a three-way ligation performed. All gene constructs were electroporated into *Agrobacterium tumefaciens* strain LBA4404 and used for genetic transformation of cassava variety NASE 13.

### Production of RNAi transgenic NASE 13 cassava plants

Cassava genetic transformation was performed following Segatto *et al*. (2022). Organized embryogenic structures (OES) were induced from leaf explants of micropropagated cv NASE 13 cultured on Murashige and Skoog basal medium (Murashige and Skoog, 1962) supplemented with 20 g/l sucrose (MS2) and 50 μM picloram. Friable embryogenic callus (FEC) was generated from OES by sequential culture on Gresshoff and Doy medium (Gresshoff and Doy, 1974) containing 20 g/l sucrose and 50 μM picloram (GD2 50P). *A. tumefaciens* strain LBA4404 carrying the binary vector of interest was grown to OD□□□ 0.6 and used to infect FEC for 45 min in medium supplemented with 200 µM acetosyringone. After four days co-culture, FECs were transferred to resting medium consisting of GD2 50P supplemented with 125 mg/l cefotaxime and cultured for 14 days. Transgenic callus tissues were recovered by subsequent subculture to GD2 50P medium containing 125 mg/l cefotaxime and 25 μM paramomycin. Growing callus colonies was transferred to MS2 medium supplemented with 5 μM NAA to induce development of mature embryos, after which cotyledon-stage embryos were transferred to MS2 containing 2 µM 6-benzylaminopurine (BAP) to induce germination. Regenerated plantlets were micropropagated and maintained on MS2 medium (Segatto *et al*., 2022).

### Production and visual assessment of plants transgenic for the GUS reporter

The p8670 binary vector was produced by inserting the *AtSUC2* promoter and *uid*A visual reporter gene into the modified pCAMBIA 2300 backbone which carries the *nptII* selectable marker gene under control of the 35S promoter (Figure S1B). This construct was used to transform FEC of cassava cv. 60444 (Segatto et al. 2022). Leaves from regenerated *in-vitro* plants, plus hand cut sections of stem and storage roots were obtained from 12-week-old greenhouse-grown plants and used for GUS assays following Jefferson *et al*., (1987), with modifications. Tissues were incubated in X-Gluc solution (1 mM X-Gluc, 100 mM sodium phosphate buffer pH 7.0, 10 mM EDTA, 0.5 mM potassium ferricyanide, 0.5 mM potassium ferrocyanide, and 0.1% Triton X-100) at 37 °C for 12 h, then cleared in 70% ethanol. Sections were mounted on glass slides and imaged using a Nikon SMZ1500 fluorescence microscope and QImaging Retiga 1300 digital camera.

### Plant establishment and growth in the greenhouse

Micropropagated *in-vitro* plants were transferred to 7.6-cm pots containing potting soil (Conrad Fafard, Agawam, MA, USA) (Segatto *et al*., 2022). After acclimatization on a mist bench, plants were grown on the open greenhouse bench under conditions of 28 °C/27 °C day/night temperatures and 70–95% relative humidity. Plants were watered with reverse-osmosis water twice times daily, and fertilized twice weekly with Jack’s professional fertilizer at 100 µg/g dry weight.

### Determination of the construct-specific RNA expression in RNAi transgenic plants

Transgenic in vitro plantlets were characterized for construct-specific RNA expression by quantitative real-time PCR (qRT-PCR) as described previously (Narayanan *et al*., 2021), using the primer pairs listed in Table S3. Total RNA was isolated using the Fruitmate kit (Fruitmate^TM^, Takara, Shiga, Japan) per manufacturer instructions, followed by on-column DNAse I treatment (Sigma Aldrich, St. Louis, MO, USA). One microgram of total RNA was reverse transcribed using the SuperScript® III-First-Strand Synthesis System (Invitrogen, Waltham, MA, USA). RNA expression in transgenic and non-transgenic control plants was quantified by qRT-PCR using the primer pairs listed in Table S3, plus 10 ng reverse transcriptase template and SsoAdvanced^TM^ Universal SYBR® Green Supermix (Bio-Rad laboratories Inc., Hercules, CA). The endogenous cassava gene *PP2A* was used as an internal control (Moreno *et al*., 2011). qRT-PCR cycling conditions included an initial denaturation holding stage at 95 °C for 30 s, followed by 39 cycles at 95 °C for 5 s, 61 °C for 30 s, and a melt curve stage from 65 °C to 95 °C with 0.5 °C increments for 5 s, followed by final extension at 95 °C for 5 min. For each sample, reactions were set up in triplicates to ensure reproducibility. Quantification of the relative transcript levels was performed using the comparative C_T_ (threshold cycle) method (Livak *et al*., 2001). To determine construct-specific RNA expression in insect-challenged transgenic plants, covered top leaves (protected from insect infestation) from transgenic and control plants subjected to insect challenge for 96 hours were collected. RNA extraction and PCR procedures were carried out using the same methodology described above.

### Detection of transgenically-derived siRNAs by Northern blot analysis

The accumulation of transgenically-derived siRNAs in transgenic plants was assessed using Northern blot analysis. Total RNA was isolated from 100 mg of fresh leaf tissue collected from *in-vitro* grown plantlets using TRIzol reagent (Ambion, Houston, TX, USA). Northern blotting, hybridization, probe preparation, and siRNA detection were carried out as described by Beyene *et al*., (2017). Thirty micrograms of total RNA were resolved on 15% Criterion™ TBE-Urea Precast Gels (Bio-Rad, Hercules, CA, USA) and transferred to Hybond™ N+ membrane (GE Healthcare Ltd) using a semi-dry transfer apparatus (Bio-Rad, Hercules, CA, USA). DIG-labeled RNA probes were synthesized using the DIG RNA Labeling Kit and SP6/T7 polymerase, following the manufacturer’s protocol (Roche Applied Science, Indianapolis, IN, USA). Signal intensity was visualized by scanning the membranes and quantified using ImageJ software version 10.2 (Rasband, 1997–2019).

### Rearing experimental whitefly colonies on eggplants

Seeds of eggplant *Solanum melongena* var. Black Beauty (Kings Seeds, UK) were sown into a 1:1 mixture of loam-based compost (J. Arthur Bower’s John Innes No 2, UK) and coir/perlite 70/30 mix compost (Jiffy Tref propagation compost, UK) in 5x5x8 cm disposable plastic pots. The plants were enclosed in a Bug Dorm cage that was thrips- and mites-proof and maintained in a whitefly-free room at 28 ± 2 °C, 50–60% relative humidity and a 14:10 light:dark photoperiod. The plants were watered twice a week with a nutrient solution containing 18:18:18+3 NPK + magnesium + trace elements (<u>1:1:1 | Products | Solufeed</u>). After four to six weeks, when the plants had at least four true leaves, adult whiteflies were released on the eggplant plantlets. The *B. tabaci* colonies from sub-Saharan Africa subgroup 1 (SSA1-SG1) used in this study were established from field populations collected from cassava in Uganda and Tanzania in 2013 (Mugerwa et al., 2021c). The purity of *B. tabaci* colonies was checked by sequencing the partial mtCO1 gene amplified using primer pairs C1-J-2195 and TL2-N-3014 (Dinsdale *et al*., 2010; Vyskočilová *et al*., 2018).

### Growing transgenic cassava plants for whole plant bioassays

Transgenic and non-transgenic plants of NASE 13 were imported from the Donald Danforth Plant Science Center (DDPSC), St Louis, MO as *in-vitro* plantlets growing in Petri dishes on MS2 medium solidified with gelzan. Plants were placed in a growth chamber maintained at 28 ± 2 °C, 50–60% relative humidity and a 14:10 Light:Dark photoperiod. After 5-7 days, plantlets were removed from the growth medium and the roots washed using lukewarm tap water to remove all the gelling agent. The plantlets were transferred to 1:1 mixture of loam-based compost (J. Arthur Bower’s John Innes No 2, UK) and coir/perlite 70/30 mix compost (Jiffy Tref propagation compost, UK) in 5x5x8 cm disposable plastic pots and placed in incubation trays with transparent lids to minimize water loss. After about eight weeks, plants with more than three or four leaves were transferred and enclosed in Lock & Lock whitefly-proof cages at one plant per each cage (Wang et al., 2011). The Lock & Lock cages had two side openings in the upper container covered by 160-µm nylon mesh. Plants were watered twice a week with a nutrient solution containing 18:18:18+3 NPK + magnesium + trace elements (<u>1:1:1 | Products | Solufeed</u>).

### Whole plant adult and nymph insect performance bioassays

The laboratory plant bioassays were performed on cassava plants enclosed in Lock & Lock whitefly-proof cages using a *B. tabaci* SSA1-SG1 colony reared on eggplants. For adult survival assays, adult whiteflies were released onto cassava plants growing in Lock-Lock cages within the NRI insectary at 28 ± 2 °C; 14:10 h light:dark cycle and 50-60 % relative humidity. Six clonal replicate plants per transgenic event were each inoculated with fifteen pairs (male and female) of 1–2-day-old, whitefly adults. Survival was recorded on the 7^th^ day after release of the insects. In the nymph development assays, fifteen pairs (male and female) of 1–2-day-old whitefly adults were released onto six-week-old cassava plants in Lock-Lock cages within the NRI insectary (28 ± 2 °C; 14:10 h light:dark cycle at 50-60 % relative humidity) for a 24 h egg laying period. Six plants (replications) per transgenic event/line were tested. After 25 days, the developmental status of the progeny was determined for number of 2nd and 3rd nymphs, number of red-eyed 4th nymphs (pupae) and number of individuals that completed development and emerged as adults, leaving the remains of their exoskeleton (exuviae) behind.

### Statistical analysis

The significance of the differences between the proportion of adult survival and the proportion of progeny that reached advanced development stages (the proportion of red-eyed 4th nymphs and exuviae from the total number of progeny) was tested using a one-way ANOVA model followed by pairwise comparisons. Prior to the analysis, the proportional data were arcsin-square root transformed. As multiple tests were conducted, a false discovery rate (FDR) correction was applied. Statistical significance was assumed at *P* ≤ 0.05. All statistical analyses were conducted using JMP Pro 18.0 (SAS Institute, Cary, NC).

### Analyses of target gene expression in whiteflies

Downregulation of target insect genes was determined from insects after feeding on RNAi transgenic, and non-transgenic control plants. Samples consisting of 50 surviving adults (after 7 days of feeding) or 50 nymphs (after 15 days of development), were collected into 1.5 ml Eppendorf tubes, immediately placed into liquid nitrogen, and stored at -80. Total RNA was extracted using an ISOLATE II RNA mini kit (Meridian Bioscience, Ohio) following the manufacturer’s protocol, with DNA contamination removed using a PerfeCTa DNaseI (Quanta Bioscience, Massachusetts) treatment. Reverse transcriptase reaction was performed using 500 ng RNA from each sample and the Verso cDNA synthesis kit with Oligo-dT primer (Thermo Fisher Scientific, Massachusetts). The expression level of each target gene was examined using qRT-PCR. The reactions were performed using the CFX Connect Real-Time PCR System (BIO-RAD, California). A set of primers (Table S3) was designed and calibrated for each target gene based on the Thornton and Basu, (2011) protocol using the Primer3 software (Kõressaar *et al*., 2018; https://primer3.ut.ee/[MOU1]). The *B. tabaci ribosomal protein L13a* was used as the reference gene. The qRT-PCR master mix contained: 5 µl iTaq Universal SYBR Green Supermix (BIO-RAD, California), 0.5 µl of both forward and reverse primers (2 pmol/µl), 2 µl DDW and 2 µl cDNA template. The expression of target and reference genes was tested in triplicate for each sample to ensure the validity of the results. qRT-PCR thermal conditions consisted of one cycle of 95 for 2 min, followed by 40 cycles of 95 for 5 s and 60 for 30 s, and an ending cycle of 95 for 5 s, 65 for 5 s, and 95 for 30 s. Quantification of the transcript’s expression levels was conducted according to the ΔΔCt method. A one-way ANOVA model was used to determine the significance of the differences between the expression means of the treatment and control samples (*P* ≤ 0.05). As multiple tests were conducted, a false discovery rate (FDR) correction was applied.

### Population dynamics modelling

#### Model – single population

The life cycle of *B, tabaci* SSA1-SG1 was modeled with four stages: eggs, nymphs, pupae and adults. Each day, individuals were considered to either remain in their current stage, progress to the next stage or die. Adults could also lay eggs. Transition probabilities depend on temperature, *T*. To simulate the dynamics, a matrix population model was used (Bodine *et al*., 2014). The population state was a 4×1 vector giving the numbers of eggs, nymphs, pupae and adults, and daily transitions, given by a 4×4 Leslie matrix, *M*(*T*), whose entries are temperature-dependent transition probabilities. Each day the current population vector was multiplied by the temperature-specific matrix. To avoid unrealistic exponential growth and to reflect a carrying capacity of ∼500 adults per leaf, density-dependent adult mortality was applied: with each day, an amount of γ× (adult population)^2^ subtracted from the adult population.

#### Parameterization – single population

The transition probabilities for the wild-type population were obtained based on laboratory measurements of development, reproduction, and mortality rates (Aregbesola *et al*., 2020). The laboratory data provided the number of individuals, out of a given starting number, that developed to the next stage, along with their median development time in days, *m_i_*, where *i* is the stage index. For our model, the daily development rates were calculated as r*_i_* =1/*m_i_* Daily mortality rate was calculated as *d_i_* = 1–survival rates at stage *i*. It follows that the fraction of population remaining in their present state is given by 1-*r_i_* - *d_i_*. Four life stages (eggs, nymphs, pupae and adults) were used, although the laboratory experiments separated the early nymph stages into two (2nd and 3rd nymphs). The nymph stages were combined into one stage by using the following calculation: *T_j_* (*j* = 1,2,3) represents the median development time from the *j*^th^ to (*j* + 1)^th^ instar nymph, and T_4_ representing the median development time required for individuals in the 4^th^ instar nymph stage to develop into pupae. The daily development rate for the entire nymph stage is given by *r*_2_ = (*T*_1_ + *T*_2_ + *T*_3_ + *T*_4_)^-1^. Accordingly, stage-specific mortality was denoted as *N_j_*, and the daily mortality rate of nymphs was calculated as *d*_2_ = (*T*_1_*N*_1_ + *T*_2_*N*_2_ + *T*_3_*N*_3_ + *T*_4_*N*_4_)/(*T*_1_ + *T*_2_ + *T*_3_ + *T*_4_). The detailed calculation of the Matrices is given in the Supplementary Data File. The daily survival rate for the nymphs was calculated as s_2_ =1 - r_2_ - d_2_.

Lab-derived parameters were adjusted to reflect field mortality using parameter q. This adjustment reduced survival rates in the egg, nymph, and pupae stages to match observed field adult emergence rates of approximately 10% (Zidon *et al*., 2016). The population dynamics under two scenarios was compared: with no intervention (wild-type plants), and with intervention (transgenic plants). The rates of the transgenic population were adjusted based on our empirical findings. Based on the findings that a fraction *P_L_* of the nymphs and *P_A_* of the adults are eliminated in a given generation. Accordingly, we increased the daily mortality rates such that an additional fraction of 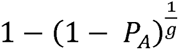 of adults and 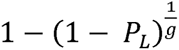 of nymphs die each day in transgenic populations, where *g* is the number of days in a generation.

The initial model included five temperatures: 16 °C, 20 °C, 24 °C, 28 °C, and 32 °C. Field measurements of temperatures in Kampala, Uganda were used, recorded at 15-minute intervals (Macdonald *et al*., 2018), and averaged to daily values. When the daily average temperature *T* fell between two of these values, interpolation using a fractional distance between lower and upper bounds was applied to calculate the matrix for any given temperature when running the simulation with actual temperatures.

#### Model – two populations

Simulated dynamics was developed for a heterogeneous field containing plots of wild-type and transgenic plants. Migration rates represent the proportion of wild-type and transgenic plants surrounding a focal transgenic plant. The two-population model uses the same temperature-dependent transition structure but extends the 4×4 matrices to 8×8, representing the insects’ dynamics on both plant types. The upper-left 4×4 block describes stages on non-transgenic plants, covering egg, nymphal, pupal and adult stages. The lower-right 4×4 block describes the corresponding stages on transgenic plants. Migration is modelled as adult movement from non-transgenic to transgenic plants. The migration rate appears in the element coupling these two stages in the extended matrix. For non-transgenic fields, migration from transgenic sites was assumed negligible, given the large extent of non-transgenic areas relative to typical dispersal distances of *B. tabaci*, and was set to zero. Temperature-specific matrices were selected and interpolated as in the single-population model, using the same temperature data.

#### Parameterization – two populations

Two migration scenarios were modelled based on field-realistic dispersal distances: (1) where maximum migration rate was 1% per day (Byrne, 1999), and (2) where maximum migration rate was 10% per day. Migration rates were scaled to represent realistic field compositions, although the true migration rates are largely unknown.

## Results

### Transgenic GUS expression under control of the SUC promoter

Transgenic plants of cassava variety 60444 were produced expressing the GUS visual marker gene under control of the SUC2 promoter. Histochemical staining of transgenic plants (Figure 1A) revealed tissue-specific GUS expression localized to the vascular tissues in leaves (Figure 1B) and in transverse sections of stem and storage roots, particularly associated with the phloem (Figure S2).

**Figure 1.**
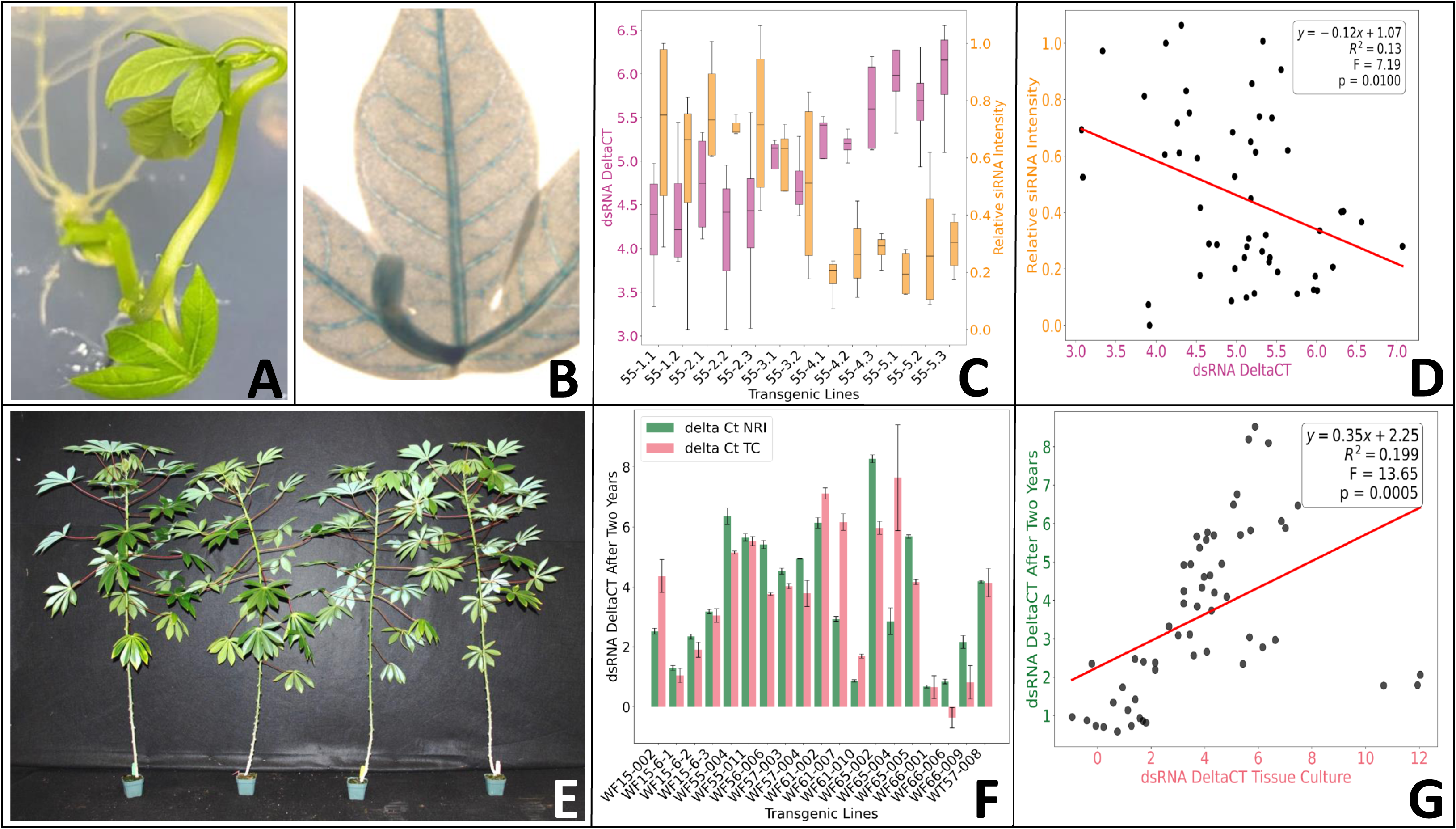
Production and analysis of insecticidal RNAi transgenic plants. (A) Transgenic plant in tissue culture. (B) The SUC2 promoter drives GUS expression to the vascular tissues of the leaves. (C) NASE 13 transgenic RNAi plants express both dsRNA and siRNA. (D) Significant positive correlation between dsRNA and siRNA levels in NASE 13 transgenic RNAi plants (higher delta Ct values = higher expression). (E) 12 weeks old NASE 13 transgenic RNAi plants display normal phenotype. (F) Stable expression of dsRNA in NASE 13 transgenic RNAi plants over two years (delta Ct NRI = two years old plants; delta Ct TC = young tissue culture plants). (G) Significant positive correlation between expression levels of dsRNA constructs over two years.

### Production and analysis of insecticidal RNAi transgenic plants in cultivar NASE 13

Fifteen RNAi gene constructs targeting the four functional strategies: osmoregulation, carbohydrate homeostasis, symbiosis and detoxification under control of the AtSUC2 promoter, plus an RNAi GFP control construct, were produced and transformed into cassava variety NASE 13. A total of 140 independent RNAi transgenic events were recovered, with the minimum target of five independent events achieved for all constructs except DWF 64 (DapF) (Table S2). Transgenic plant lines were evaluated for their ability to express RNA from the inverted repeat sequence using qRT-PCR (Figure 1C). To show that template was being processed into siRNAs, selected plants were also analysed for accumulation of small interfering RNA (siRNA) species specific to the inverted repeat sequence (Figure 1C). A clear positive correlation (higher delta Ct values reflects higher expression) was observed between dsRNA and siRNA expression levels (Figure 1D). The transgenic plant lines were screened under greenhouse conditions, where no adverse phenotypic effects were observed associated with expression of the inverted repeat sequences (Figure 1E). Stability of transgenic dsRNA expression was also assessed after a two-year period (Figure 1F). These longitudinal analyses demonstrated sustained expression of the dsRNA constructs and a consistent correlation of expression levels across the two-year timeframe (Figure 1G).

### Insecticidal effects of the four strategies on whitefly performance in whole plant bioassays and their correlation with gene silencing

Transgenic NASE 13 cassava plants expressing the 15 different RNAi constructs targeting genes within the four strategies underwent whole plant bioassays to assess their insecticidal effect on adult and nymph whiteflies. The most insecticidal transgenic lines in each strategy caused up to 35-58% decrease in adult survival after seven days feeding when compared to the non-transgenic wildtype (WT) control line NASE 13. Excluding the two constructs DWF59 and DWF60, at least two transgenic events, showed significant (*P* ≤ 0.05) negative effect (reduced adult survival) when compared to an event expressing dsRNA against the GFP gene (Figure 2A). Of special interest were constructs DWF55, expressing dsRNA targeting an *ST* gene; DWF56, expressing dsRNA targeting an *ABC* transporter gene; DWF61, expressing dsRNA targeting the *ArgH* gene; DWF63, expressing dsRNA targeting the *LysA* gene and DWF66, expressing dsRNA targeting the *UGP* gene. When using these constructs, 83-100% of the transformed plant lines were capable of causing a significant reduction (*P* ≤ 0.05) in adult survival, indicating a strong negative effect of the specific targeting RNAi. In contrast, only two transgenic events, DWF66-006 and DWF63-001, out of a total of 104, presented high efficacy capable of reducing the adult whitefly population by more than 50%.

**Figure 2.**
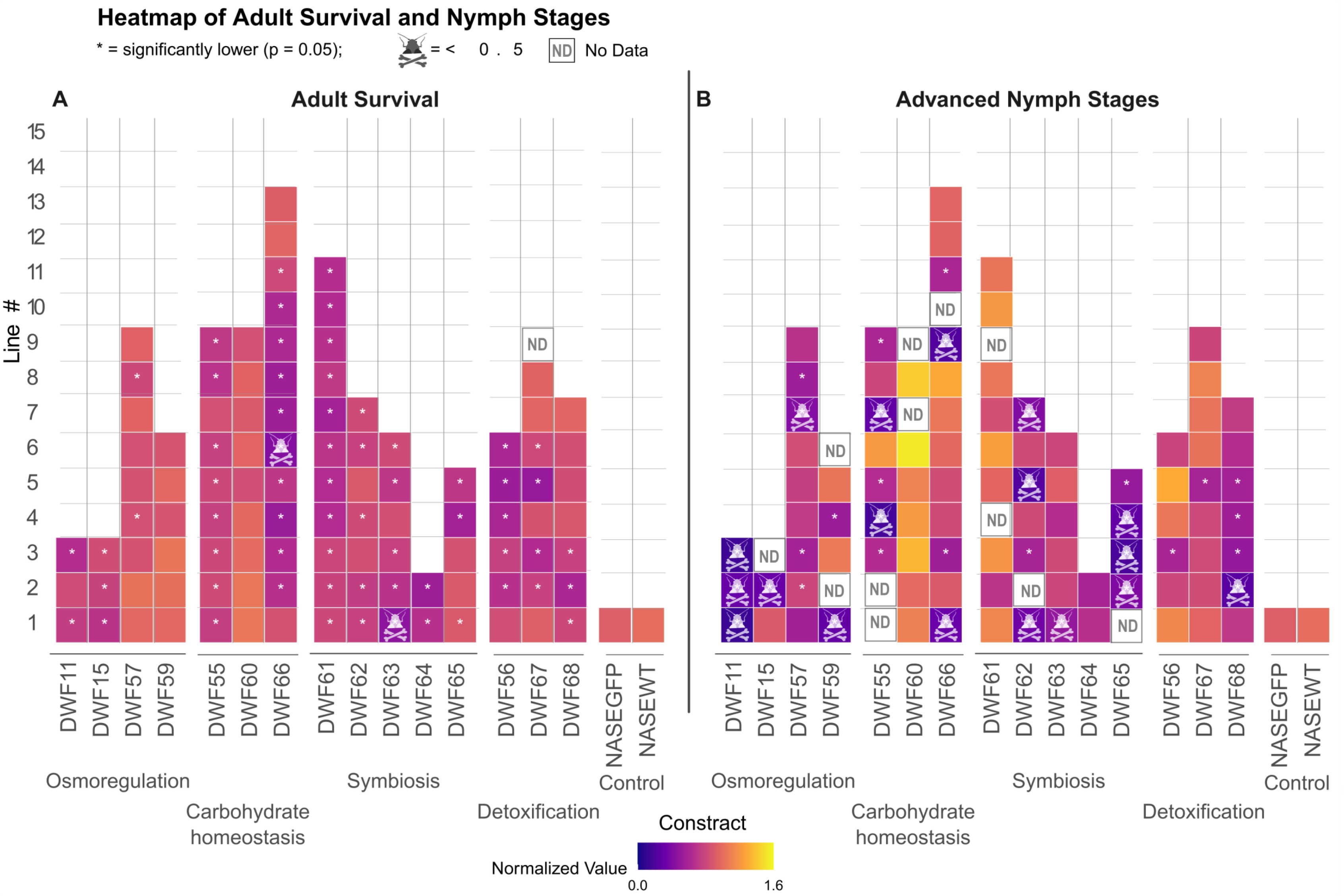
Insecticidal effects of the four strategies (osmoregulation, carbohydrate homeostasis, symbiosis and detoxification) on whitefly performance in whole plant bioassays. (A) The proportion of adults that survived on NASE 13 transgenic RNAi plants. Data was normalized to the mean survival proportion of the control (NASE WT plants). Excluding the DWF59 and DWF60 constructs, at least two lines showed significant (* = *P* ≤ 0.05) negative effect (reduced adult survival) when compared to a NASE 13 line expressing dsRNA against the GFP gene (NASEGFP). Only two transgenic events (DWF66-006 and DWF63-001) presented high efficacy capable of reducing the adult whitefly population by more than 50%. ND = no data available. (B) The proportion of progeny that reached advanced development stages (red-eye 4th nymphs and newly emerged adults) 25 days after egg laying. Data was normalized to the mean survival proportion of the control (NASEWT plants). Eight of the 15 constructs showed significant (* = *P* ≤ 0.05) effect (reduced nymph development), in at least two lines, when compared to the control expressing RNAi against GFP (NASEGFP). Transformations with the DWF11, DWF55, DWF62 and DWF65 constructs produced at least two or more independent lines that presented high efficacy capable of reducing the development rate of the immature stages by more than 50%. ND = no data available. Significance was tested using a one-way ANOVA model followed by pairwise comparisons and FDR correction.

In the nymph development assays, the best transgenic events in each strategy caused 75-90% delay in development when the normalized proportion of progeny in advanced stages (the proportion of red-eyed 4^th^ nymphs and nymphs that completed development) was compared to the NASE 13 WT control (Figure 2B). Eight of the 15 constructs showed significant (*P* ≤ 0.05) effect (reduced nymph development), in at least two independent events, when compared to the control expressing RNAi against GFP (Figure 2B). Of special interest were constructs DWF11, expressing dsRNA targeting an *AQP* gene; DWF55, expressing dsRNA targeting an *ST* gene; DWF62, expressing dsRNA targeting the *CM* gene; and DWF65, expressing dsRNA targeting the *BioB* gene. When using these constructs for transformation, 68-100% of the transgenic events were capable of causing a significant (*P* ≤ 0.05) delay in nymph development, indicating a strong negative effect of the specific RNAi on immature whitefly stages. Transformations with all four constructs produced two or more independent events that presented high insecticidal efficacy capable of reducing the development of immature stages by more than 50% (Figure 2B).

We employed qRT-PCR to establish a direct association between downregulation of target gene expression and reduced performance of whitefly nymphs and adults. RNA was extracted from nymphs and adults feeding on transgenic cassava plant lines expressing RNAi inverted repeat constructs DWF55, DWF56, DWF57, DWF61, DWF65, DWF66, and DWF68, and from control plants expressing dsGFP. In adult whiteflies, target gene expression was reduced by up to 45% relative to control adults, while in nymphs, downregulation reached 57% compared to controls. A significant positive correlation (*P*=0.0117) was observed between the level of target gene silencing and the insecticidal effect on nymph development and adult survival (Figure 3), providing proof-of-principle that suppression of target gene expression is causally linked to reduced performance in both immature and adult stages of cassava-whitefly.

**Figure 3.**
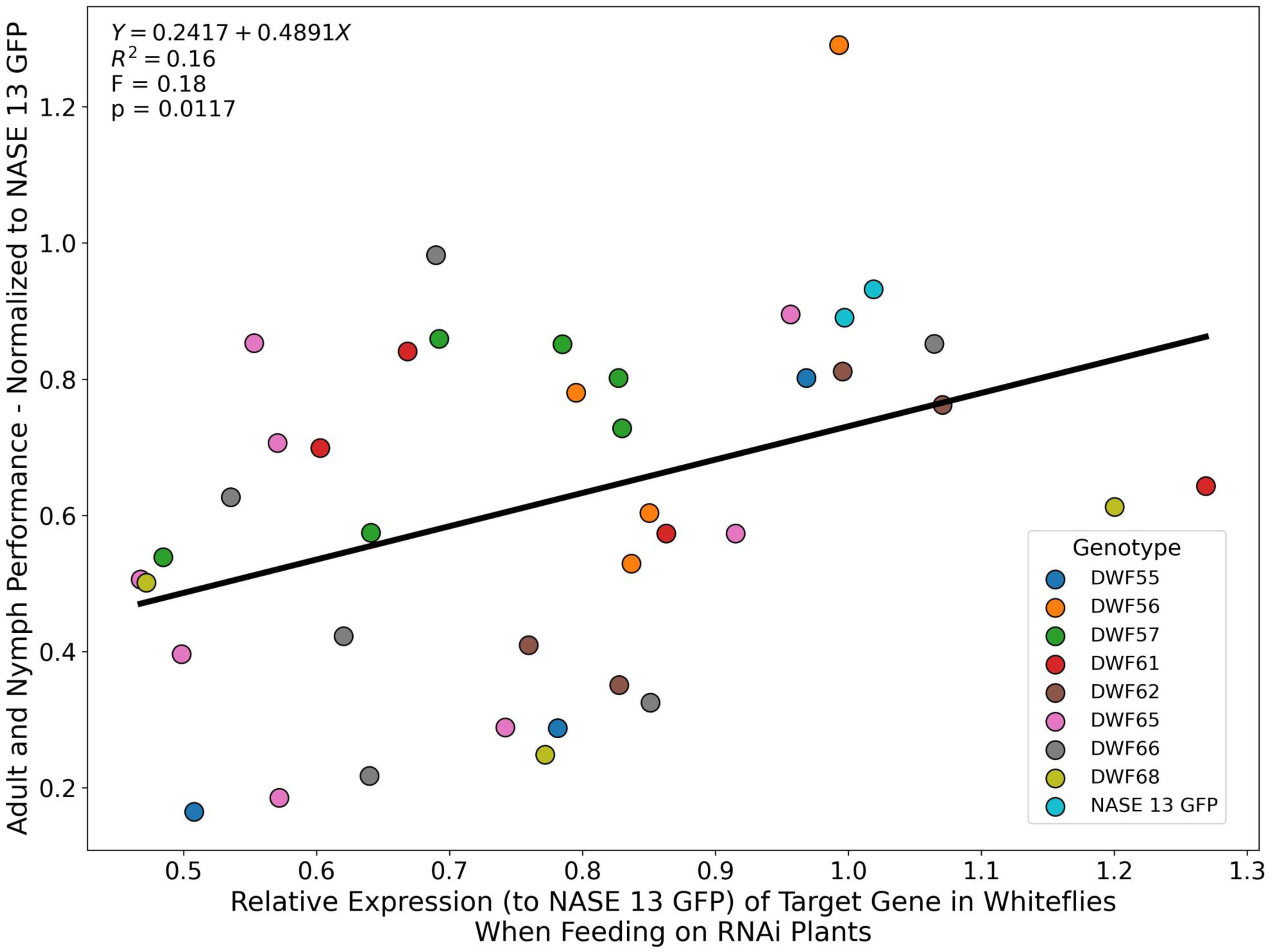
The correlation between the relative expression levels of genes targeted by the dsRNA treatments and the performance of the insects (adult survival after 7 days and nymph development stage after 25 days) after normalization to the control (NASE 13 plants expressing dsGFP). Significant positive correlation indicates that reduced gene expression results in reduced performance, indicating that the gene silencing is likely to be the mechanism causing lower performance.

### Assessing the influence of RNAi transgenic cassava on whitefly population dynamics using a temperature-driven model

Using a daily, temperature-dependent 4×4 Leslie matrix that captures transitions among egg, nymph, pupa, and adult - parameterized from laboratory measurements across 16–32 °C and driven by field temperature time series via interpolation, we simulated stage-specific *B. tabaci* SSA1 whitefly mortality of adults, nymphs, or both. Figure 4 illustrates modelled whitefly populations over the course of one year, excluding immigration and emigration processes. Simulation results indicated that population growth begins approximately 40 days after initialization, corresponding to two generations, in non-transgenic cassava fields with an exponential increase in population size occurring over the subsequent ∼40 days (two additional generations), followed by stabilization at a steady-state level after four generations. In transgenic scenarios, RNAi cassava plants were simulated to reduce adult survival (Figure 4A), nymphal development (Figure 4B), or both (Figure 4C) by 30–80%. These interventions altered population dynamics in two principal ways. First, the initial population build-up was delayed by one to five generations when the combined negative effect on both nymph and adult stages reached 60%, 70%, or 80%, respectively. Second, the steady-state population size was reduced by approximately 33%, 44%, and 68% under the same respective treatment intensities. Two additional insights can be drawn from the simulations presented in Figure 4. First, RNAi technologies targeting immature nymphal stages are predicted to be more effective under field conditions than those primarily affecting adult stages. Second, interventions with a combined negative effect below 50–60% are projected to exert only minor influence on the overall population dynamics of whiteflies in the field.

**Figure 4.**
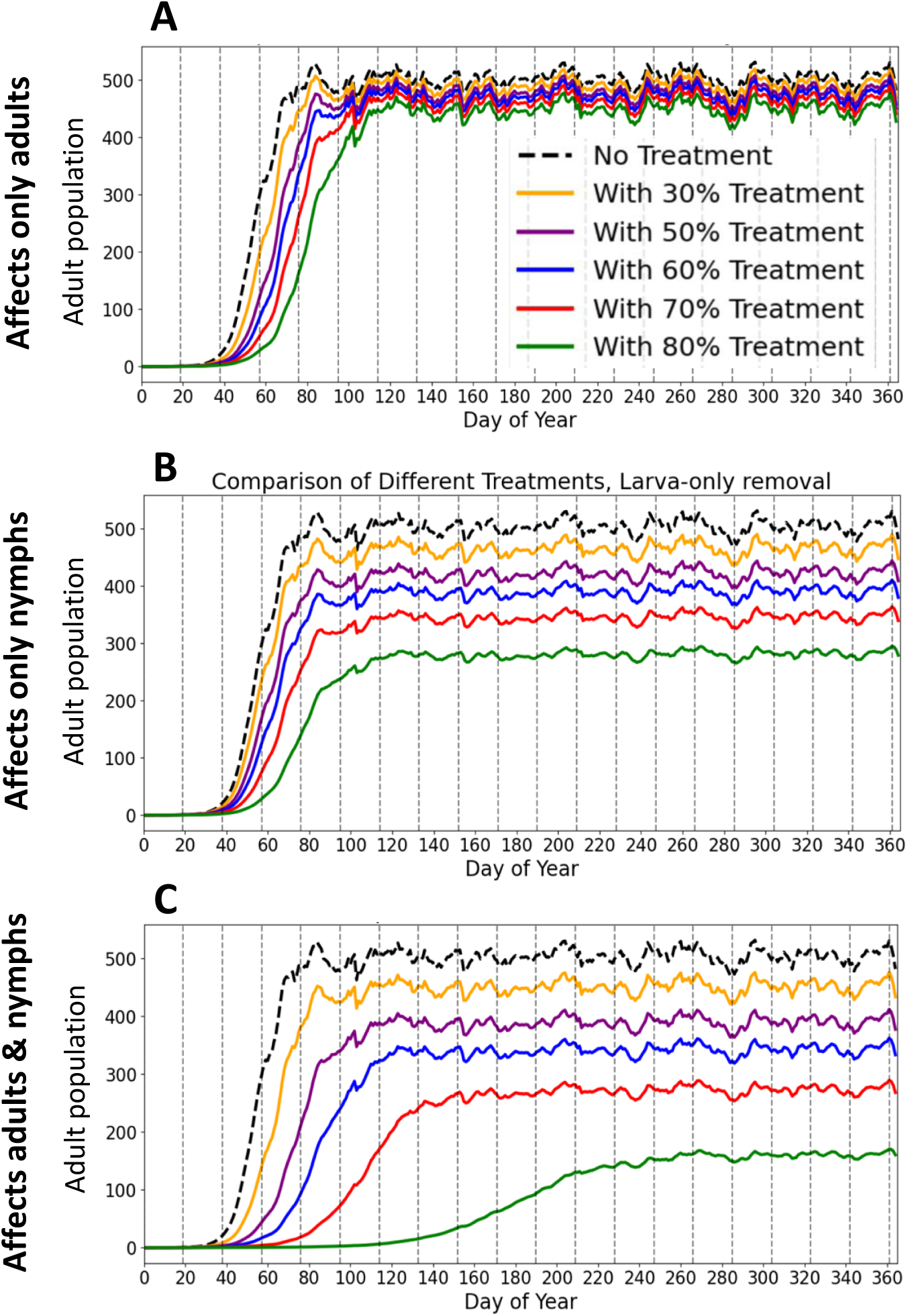
Adult population density over time in transgenic cassava. Colours represent the percentage reduction applied to adults and/or nymphs, with each curve giving the adult population trajectory. Both the strength of reduction (colours) and the affected life stage (panels) are examined. (A) both adults and nymphs are affected, (B) only nymphs are affected, and (C) only adults are affected. Higher percentages of reduction make the adult population grow more slowly and stabilize at a lower level. Nymph treatment has the greatest impact: adult-only treatment is largely ineffective, nymph-only treatment substantially reduces adult density, and affecting both stages is most effective.

We extended the model to two interacting plots (transgenic and non-transgenic) by doubling the state space (8×8) and allowing daily adult migration between plots, and then varied the fraction of the landscape planted with transgenic cassava while comparing 1% versus 10% daily movement. A constant 60% reduction in both nymphal development and adult survival was assumed for the transgenic plants. Simulation results showed that at a 1% migration rate, population growth dynamics exhibited only minor differences among the four mixed-field scenarios (Figure 5A). However, when the migration rate was increased to 10%, substantial differences emerged in whitefly population dynamics between field mixtures dominated by transgenic plants and those dominated by non-transgenic plants (Figure 5B). Given the absence of reliable estimates for adult SSA1 whitefly migration rates in the current literature, we conservatively limited the migration parameter to a maximum of 10%. Nonetheless, the simulation outcomes suggest that migration rates exceeding 10% could significantly compromise the effectiveness of RNAi transgenic cassava in agricultural landscapes where non-transgenic, whitefly-susceptible varieties remain prevalent.

**Figure 5:**
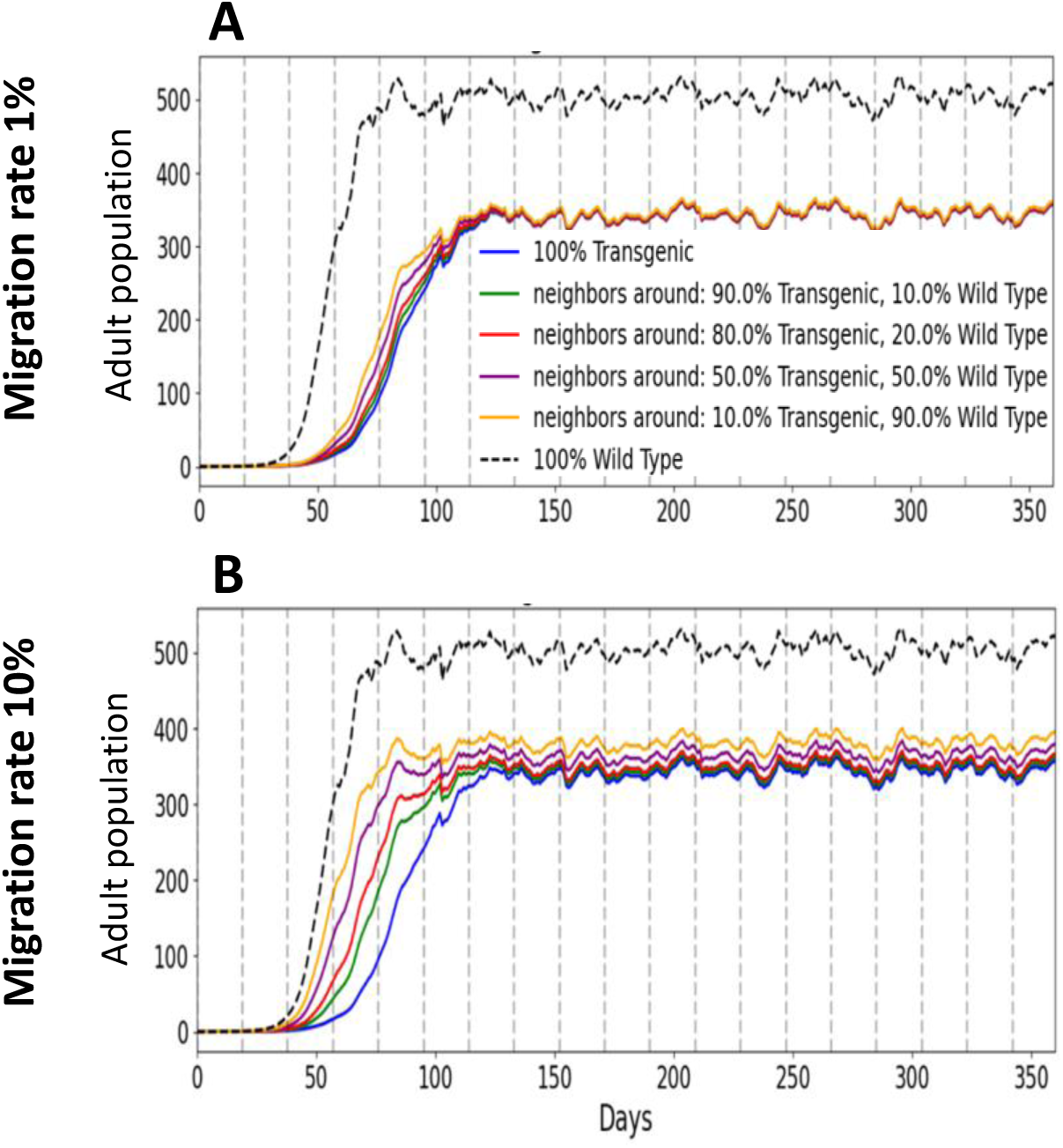
Adult population density over time for a transgenic plot in the presence of a neighbouring non-transgenic plot. Colours indicate the percent of total area planted with transgenic cassava; each curve shows the adult population density over time for that coverage level. We vary both coverage (colours) and daily movement between plots: (A) 1% migration per day; (B) 10% migration per day. Greater transgenic coverage slows growth and lowers the final level. At the same time, transgenic cassava remains effective even with migration from the non-transgenic plot, even at 10% daily migration rate.

## Discussion

We report development of efficacious *in planta* RNAi technology for controlling a major insect pest of African cassava, the cassava-whitefly *B. tabaci*. Cassava is a strategic food security crop for smallholder farmers, particularly in sub-Saharan Africa (Obong’o *et al*., 2024). Integrated management packages of the whitefly vector are currently limited by the relatively low availability of improved disease or insect resistant varieties, and the fact that commercial insecticides and other input-intensive measures are expensive and/or logistically inaccessible for the majority of smallholder farmers (Legg *et al*., 2022, Bayiyana *et al*., 2023).

We bring evidence that transgenic cassava plants in the East Africa cultivar NASE 13, expressing RNAi technology that targets essential genes in whitefly biology, has potential to significantly supress whitefly populations. The most efficacious transgenic events in each of the four reported strategies showed moderate ability (35-58%) to decrease adult survival, but were highly effective (75-90%) in delaying nymph development. In this context, it is important to note that observations during the bioassay indicated that in the majority of cases, developmental delay caused by RNAi prevented successful completion of the insect life cycle, eventually leading to nymph death. The potential of the RNAi technology to achieve significant population suppression of cassava whiteflies was emphasized by two complementing outcomes of the simulation models. First, whitefly control strategies targeting immature nymphal stages were predicted to be more effective under field conditions than those primarily affecting adult stages. Second, only control strategies with a proven potential to cause at least 60% reduction in nymph performance should be taken into consideration as possible new strategies for efficient whitefly control. Our simulation models also explored the important effect of adult migration from fields planted with whitefly susceptible, non-RNAi cassava plants, a topic that is neglected in most RNAi studies to date. The simulations indicated that the technology will be most effective if applied at a regional level with high percentage of farmers growing the RNAi varieties.

In smallholder farming systems, such as that of most cassava cultivation in Africa, the main advantage of i*n planta* dsRNA expression is its ability to provide sustainable, seasonal-long, transgenerational protection against whiteflies with no farmer-required inputs. Moreover, once an efficacious RNAi expression cassette is developed it can be introduced into any farmer-preferred variety through direct transformation or conventional backcrossing (Basso *et al*., 2025). A growing body of studies indicate that RNAi-based insect control is highly sequence-specific and degrades rapidly in soil and water, so that, when target genes are selected using bioinformatic filters to minimize homology with non-targets, the probability of ecologically relevant off-target effects is low to the point of negligible, compared with the negative effects of broad-spectrum chemical insecticides (Chen and Schutter, 2024). Empirical non-target studies, including recent assessments of dsRNA-expressing cotton on the predatory ladybird *Harmonia axyridis* (Yao *et al*., 2025) and dietary exposure assays with multiple beneficial insects, generally report no adverse effects on survival, development or fitness under realistic exposure scenarios (Castellanos *et al*., 2022; Taning *et al*., 2021). Finally, *in-planta* dsRNA expression for pest control is advantageous over repeated dsRNA foliar spraying in marginally profitable and vulnerable crop production systems such as smallholder farms. This is mainly because dsRNA spraying requires assess to, and involves recurrent costs of purchasing and applying the formulated dsRNA that are not feasible for small holder farmers (Bramlett *et al*., 2020; Willow and Veromann, 2022).

There are limitations to the RNAi technology that should be taken into consideration to ensure durable, long-term efficacy. A central concern is the evolution of resistance in target pest populations (Palli, 2023). Two possible scenarios should be considered in this respect. In the first, resistance is specific to a particular RNAi target (e.g. changes in the target gene sequence or its regulation), while the core RNAi machinery remains functional. In this case, resistance management can simply include switching to an alternative sequence region within the same gene or to a different essential gene to restore susceptibility (Hashiro and Yasueda, 2022). In the second scenario, insects evolve resistance that broadly impairs the RNAi pathway itself. For example, increased dsRNA degradation in the insect gut that prevents effective gene silencing (Spit *et al*., 2017) or reduced dsRNA uptake by the insect, often associated with downregulation or mutations in dsRNA transporters such as Sid-1-like proteins or clathrin-mediated endocytosis components (Duanmu *et al*., 2025; Khajuria *et al*., 2018). To overcome the second scenario, resistance management (IRM) frameworks for dsRNA traits recommend pyramiding (stacking) with at least one additional technology that operates through a different insecticidal mode of action (Narva *et al*., 2025). In this case, only insects that will evolve resistance to both technologies will survive, which is less likely to happen (Ni *et al*., 2017). If a second technology is not available, the time until resistance to the RNAi technology evolves can be significantly extended by maintaining non-RNAi refuges that produce large numbers of susceptible insects that dilute the resistance alleles (Narva *et al*., 2025). This approach was successfully applied during the commercialization of the *Bacillus thuringiensis* (Bt)-derived technology, especially if the evolved resistance is associated with a significant fitness cost (Carrière and Tabashnik, 2023).

## Conclusion

Evidence provided here indicates that *in-planta* RNAi technology offers a realistic option in developing durable control of cassava whitefly in African cassava fields. Our work and previous studies suggest that targeting genes involved in osmoregulation, such as aquaporin, sugar transporter and α-glucosidase, and/or symbiosis, such as chorismate mutase and biotin synthase, are good targets for markedly reducing the ability of the insect to build large populations in the field (Luo *et al*., 2017; Wang and Luan, 2023; Wintraube *et al*., 2025). A logical next step to enhance insecticidal activity is to drive higher and more spatially appropriate dsRNA accumulation by exploiting additional strong phloem-specific promoters such as MeSWEET1-like, and MeSUS1 (Zierer *et al*., 2022). In parallel, stacking dsRNAs against multiple essential whitefly genes or co-targeting dsRNases and vital genes has been proposed, as a way to increase mortality and slow resistance evolution (Luo *et al*., 2017; Nitnavare *et al*., 2021). With these improvements, all achievable in a relatively short period, plant-mediated RNAi will be well positioned to become a core, resistance-management-compatible tool for whitefly control in cassava fields.

## Acknowledgements

We thank Angela E. Douglas for contributions towards the RNAi gene constructs DWF10-15 and her insightful comments during development of the experimental system. The NASE 13 variety utilized here was provided by the National Crops Resources Research Institute, NARO, Uganda. We also thank Ksenia Juravel for her help during Figures’ production. This work was supported by the Bill & Melinda Gates Foundation, Seattle, WA, through the University of Greenwich (Grant number OPP1058938).

## Author contributions

OM designed the dsRNA constructs. EY, NN and JG produced the RNAi gene constructs. TJ produced the RNAi transgenic plants. NN and JG determined the construct-specific RNA expression in RNAi transgenic plants. OM, SM, NJT, JC and RARS designed the plant bioassays. RARS conducted the insect performance plant bioassays. DW and SL determined the target gene expression levels of RNAi treated insects. SM analysed the data. OH and AL conducted the population dynamics modelling. OM, NN, AL, NJT and SM wrote the manuscript. JC, NJT and SM obtained funding and provided other essential resources. All authors approved the final manuscript.

## Data availability statement

Modelling code and data associated with this paper (Supplementary Data File) was published in Dryad (code via Zenodo in the same link).

Link for reviewers:

http://datadryad.org/share/LINK_NOT_FOR_PUBLICATION/HlF-Xa6LHfo_g5UHR1C5ZLgLlHBub93M_tp3lDZD8Jg

DOI: https://doi.org/10.5061/dryad.z08kprrv8

Dataset name: “Data and Code for modeling RNAi-technology effect on plant resistance for Cassava Whitefly (*Bemisia tabaci*)”. Oren Hasson and Adam Lampert.

## Supporting Information

Additional supporting information may be found online in the Supporting Information section at the end of the article.

**Figure S1.** Schematic representation of gene constructs used for the genetic transformation of cassava.

**Figure S2.** Transverse sections of stem and storage roots indicating GUS localization, particularly in phloem tissues.

**Table S1.** Candidate gene sequences with the region targeted by the dsRNA constructs highlighted.

**Table S2.** Strategy, genes and their respective metabolic pathways targeted for RNAi downregulation and number of independent transgenic events produced from each inverted repeat construct.

**Table S3.** List of qRT-PCR primers used in this study.

