## Supplementary material for "Creating resistance to the whitefly *Bemisia tabaci* in cassava through RNAi-mediated targeting of multiple insect metabolic processes": Table S1

**Table S1: Candidate gene sequences with the region targeted by the dsRNA constructs highlighted in light blue.**

N indicates that the nucleotide identity in currently not determined in the databases.

**Class I**

**DWF-11,15, ENSSSA1UGG010352, SSA1-SG1 Aquaporin**

ATGGAGGACATATCATCTTCCGGCGAAGAAATCAGCATGAAAGCGATCTCAAAAGTGATTGGAGTGCCCGATATTCGAGATGGGCCCACTCTCACAAAATGCATAGTTGCTGAGTTCGTTGGGACTTTGCTGTTAGTGCTCATAGGATGCATGTCGGTAGCATTTGTCCATCAGGAAACTTCGTTGACGTTGTGAAAATTGCCATGGCTTTCGGGCTCATCATCGCCTCTATGGTCCAGGCAATAGGTCACGTTAGTGGTTGTCACATCAATCCGGCTGTAACTTGTGGACTAGCTGTGTCGGGACATGTTAGCATAATAAAAGGTATGCTGTACATTGTCGCGCAATGCCTTGGAGCCATCTGTGGAGCAATCATTTTGAATGAAATCACGCCAAAAACAGGTTACACGGCTGCTGGTAATCTCGGAGTAACGACACTGTCTACAGGAGTTTCCGACCTGCAGGGTGTGGCGATAGAGGCACTAATCACATTTGTGCTGCTTTTAGTTGTCCAGTCCGTCTGCGATGGGAAGCGGACCGACATCAAAGGATCTATCGGTGTTGCGATAGGATTCGCAATTGCTTGCTGCCATCTCGCCGCGATCAAGTACACCGGTGCTAGTATGAACCCTGCTAGATCATTAGGCCCTGCATTCGTCAGTGGAATTTGGGACAAACATTGGGTCTACTGGGCCGGTCCAATACTCGGTGGAGTCACTGCCAGTCTGCTTTACGCCATCACTTTCAAAGCCAAAAAAAGATCGGATGAGAGCTCCTATGATTTCTGA

**DWF-15, ENSSSA1UGG011937, SSA1-SG1 alpha glucosidase 13**

ATGGAAGAGCAACCGCAACGCGATCTAGCTTGGTGGGAGAAAGGAGTTATCTATCAAATCTATCCGCGATCATTCAAAGACTCGGATGGCGACGGAGTCGGCGATTTGAAAGGAATTGCGGAGAAAATTGATTACCTATCAAAACTAGGCGTCGAAGCAGTTTGGATTTCTCCGATTTTTCGCTCCCCGATGGCAGATTTTGGTTACGATATATCGGATTTCAGAGCGATCGAGCCAATGTTTGGCACAATGGAAGATTTTGAAAAGCTGAAAAGATTATTCCATAAAAATGGATTAAAAATGATTCTGGACTTCGTGCCAAACCATACCAGTGACGAACATGATTGGTTTAAGAAATCAGTCGCAAAAGTTGATCCGTACACAGATTATTACAACTGGGTCAACGGGAAGATGAGTGAGAATGGAACCATGCAACCACCGAATAACTGGCTTTCATATTTTGGTGGGTCAGCTTGGACTTGGAATGAAAAACGAAAGCAGTACTATCTGCATCAGTTCCATCCGAAACAACCAGATTTGAACTATAGGAACCCTCTCGTTGTTCAAGAGATGAAGGATGTTTTGACTTATTGGATGGACAAAGGAGTAGATGGATTCAGGATGGACGCTGTGATGACCATCATGGAGGACATTAAACTGAGGGACGAACCACTCTCAGGCAAAACGGACGTCCTGCCAACTGATGAGGAATACCTTAACCACATTTATACCAGAGACCAACCGGGGACCTACGAGATAATTAAACAGTTTCGAAAACATCTCGATGATTATTCCAGGAAAACTCACACGGTCAAGTTCATGGCGACAGAGGCATATTCGAATTTGACCAGCACGATGAAATATTACGGCACAAAAGATAATCCTGGAGCCCACTTTACGTTCAATTTTGAGACCATCGAAGCGCTAACCCCACAATCGAACGCGGCAGATTTCAAAGAGGTCATCGAAGACTGGTATGCTGCTCTCCCTGCCGGCAAGTGGTCTAATTGGGTGTTGGGAAACCACGACAAGCGGCGGGTAGGCTCGCGCTACGGGATCGACATGTTGGACGGGCTGCACATGCTTCAGATGTGCCTGCACGGTACCTCGGTGACGTATGCCGGGGACGAGATCGGAATGGTGGAACGTTCATCCGCTGGGATCAAACCAAAGATCCGCCCGCTCTGAACGTCGGCCCTGAGAGGTATCAAAGATTCACCCGGGACCCCGCCAGAACTCCGTTCCAGTGGAATGCGTCCACAAGTGCAGGTTTTTCGACGAATCCAAAAACTTGGCTTCCAGTCAATCCTGATTATTGGTCACACAATCTAGTTACAGAAAAGAAAAAGGCCAGTAGCCATTTCAAAAATTACCGAAGATTGTTGGCTCTGAAAAAGTCGCCCGTTATTCAATTTGGCAGTGTAAATGTCTACACCCTATCAGACTGGGTCCTTGTCATAACCAGAACTCTCAAAGACCACCCGACATACATTGTCATTCTGAACCTTGGCACAGAATTAGAGGATACGAAAGGGTTGAGAACTATCGCGAACTTGCCAGACCAAATCAAACTGCATACGTGCAGCATCAACTGTGGCTACACGCCTGGAGCTCAACTTCGGACGGATGAAATCCAGCTTCGACCGAAGGCCGGCTTTGTGTGCATCGCTCGGGGTGGTCTAAAAGCCTCTTGA

**DWF-57, ENSSSA1UGG000414, SSA1-SG1 alpha glucosidase 13**

ATGGAGAGCCTACTCTACTTTGTTGTGTGCGCCAACATTTGGATTGGATTTTTGGAAAGCCATCAGATTCTGAGCAGCGGTGGAATGAAGGACTGGTGGGAGAGCACTCTGATTTATCAGATATACCCGCGCTCCTTTAAGGATTCAAATGGGGATGGCTACGGTGATCTAAATGGTATTCTCGAAAGACTGGATTACGTCAAGGAGATAGGCGTGGACACAATTTGGATTCAACCGTTCTACAAGTCGCCGATGGCTGATATGGGCTACGACGTTGCTGATTATCGAGCTGTCGATCCAATGTTCGGTACTATGGATGATTTCAAGAGGTTACTCAAAAATAGTCATGAGAAAGGGATGAAAGTGATTGTTGACTTTGTGCCGAATCATAGTAGTGATCAATGTGAGTGGTTCAAGCTCTCCGAACAACGTATCGAACCTTTCACAGATTTCTACGTTTGGAAGGACGGTAAACGGATTAACGAAACTCACACAACACATCCAAATAATTGGCGAAGCATTTTCCGAGGTCCAGCATGGACTTGGAGTGAAAAAAGGAAACAATATTACTTGCATCAGTTTGCGCCGCAACAGCCGGATCTTAACTATTACAGCCCTCGTGTGCGTCGAGAAATGGAGGATGTACTGAGGTACTGGTTGGACTTGGGGGTCGACGGATTCCGAGTTGACGCAGTGAATCACTTGTTGGAGGCGGAGCACTACGCGGATGCGCCAGCCGGCACCAAAGGAGGAGGACTCGGTGTCATCGAAGGAGTGTCACTGGTTTACACGTTACGCCAACCTGGAAATTTGCAACTTATCAAATCATGGCGGGCAGTTCTTGACGAGTATGCCAAAAAGGATGGACAAGCAAGATTATTGAGCACGGAAGCTTACGATATGCCTAAAATACTGAGCCTTCAGTTGGGGAACGCCACTTACCCTGGAGCTCAAATTCCATTTTTCGTTCGGATGGCATGGCTCGATTACGATACCACCGCGTCAAAGCTGGACTCTTACATCCACGAAATGATTGAATCCTTCCCTGTTCGCTCCTTCAATTGGGCGAGGGACAACCACGACAATCCTAGGGTGAGCACGCGATACTACCCGGAGAGCGTGGAAGCGTGGAACATGCTACTTCTCCTCATGCCCGGAGCCACGTGTATTTATTACGGGAGCGAGCTCGGGATGGAAGACGTCCATGTCCGCGATAATCAACGTCGAGACCTTCAAAACGCCGGCGGTGATCGCACGGAGACTCGGGACGGATGCAGATCCCCCATGCTCTGGGATAATACCAGAAATGCAGGGTTCACGACTGCGAAAAATCCGTGGCTACCTGTCCACCCGAACTACTGGCGGACAAATGTAGAGGCTCAAAGGCAGGACCCCAACAGCCACCTCAACATTTACAAACGACTTGTTGAGCTTAGAAGGATGCCTGTCATCCGTCATGGAGATTTGCGCACCTATGTGCTGGAGGAAATGGTGTTCATGTTTACAAGGTCTTTCAAGAGTGATACTGTAGCAGTGCTGATGAATTTAGGATCAGAAACTGAGAAGATTTGCGTCCAAGATTCAGCCTCACTGTTACCGGACACGATGTTCGTTCATACCAGAAGTCCTAATTCAGGGATGAACGTTGGAGAGGAGGTGAAAATAAGAGATGCGGGCAAAACGGAATGCATTTACATGAGGCCACAATCGGGGCTTGTACTGAGCACATTTCTCAGTGAGTCGATGAGCCCAAATAGTATATCCAGTGATGCTCCACTCTCGTCGTGGTCTTTCGCTGTTCTTCTACTAGCAATTTGTTTAAAAATCTTCATCAGCTAG

**DWF-59, ENSSSA1UGG002701, SSA1-SG1 alpha glucosidase 13**

ATGTTTCTTGAAATTTCTCGGGAGGTTTTTGGTACACCAGTTTTCACACGTACAAACCATGAAGAATTGAACGATAGGAATACGATCATGAAGGCTGAGGTTGAAGGGAAAAAACAAGCAGGTCGAAAGGGGTTGACGAAGCGCTTTCTCATCACCGTGGGACTTCTTGCGGCAATCGCATTAGGACCGTCCGGATGCGAGGGATATGGCATTCCAGACATAGAAGACCCCAAGTGGTGGCAAAACATGGTTGTGTACCAGATCTACCCGCAGTCTTTCTACGATTCTGATGGAGACGGAGTCGGCGATCTAAAAGGAATCAAAGAGAAAGTCCACTATCTTTGTGAGCTGGGAGTGACCGCAATTTGATAAACCCGTTCTACGAATCCCCGCTGAAAGACAACGGTTATGACATCTCTAATTTCAAGAAAATCCACCACAAGTTCGGTAACATGTGTGACTTTAAGGCACTCATGAAAAGACTTGATAATAATCCTTGCGGTGATAACAAAGAGATCAGAGTGATCATGGATTTCGTACCCAACCACACGAGCGACGAACATGAATGGTTTCAGAAGTCAGTCGATCGGGTGCCTCCATACAATGATTACTACGTATGGATGGATCCTAAAGGATGGGACAAAAATGGAAATCCAATCCCTCCCAACAATTGGGCATCAGTCTTCAGCTACGGAGAGGAAGCGTCTGGATGGGAGTGGAATGAGAAGAGACAACAATTTTACTTTCACGCTTTCATGAGGGAACAGCCGGACCTTAACCTTCGTTTTTCTGAAGTCAGAGACGAATTGAAGGATATCTTGAAGTTTTGGATGGATATGGGAGTAGATGGGTTCCGCATGGACGCCGTGCCGCATCTCATCGAGGACCCACAATTCCGCGACGAGGACCCCTGGTGGGATCCAAATACACGCAGAACCTACCGCAAACCTACGAAGTCATCAAAGAGCTCCGAAATTTCTTCGAGGAATACGACAAAATAAACAATCGAGTCACGTACATGACTGTGGAAGCATATGCATCGCCAGATAATACTAGACGGTACTACGGCACACTAGATGCACCTGGTGCTCACTACCCATTCAACTTTCTTCTCCTCACTGATTTTAATTCGGACACGAATGCCGTCAGCCTCAAAAACTCCATTGACTTTTACTTGAGTCAACTCAACGACTTTCAGCTTCCCAACTGGTTGTCCGGGAACCACGACTGGCCGCGAATCCCATCGAGGCTGGGGGCCTCCACCTCAGTAGATTTGATCAGCATATTGAACCTGCTGTTGCCCGGATCAAGCTGCGTGTACATGGGAGACGAGATCGGCATGGACAGCTACTGGGCCATTGACAGGGACAAGTGTCCACCAGGAGCTGCCGATGACTATCGCACGCCCAGCAGAACACCCTTCCACTGGAACAAAACTCAAAACGCAGGTTTTACGACGAACGAAGACCCGTGGATTCCTGTGAACCCCGAGTATTACAGGCTGAATGTAGAGTCAGAACGCTACCCATCATGCTCGTACTGCTCCACCCACTTGACCAATTTCAAGCACTGGTCCAGCTCAAAAAGGAATATGCGTTTATGTATGGGCCTTTATCTACCTACACTCTCACCGATGATGTCTTCTGTTTTACCAGGGAGGATTCAGAAACGATATATGTGACAGTCATGAACGTGGGTACGAATTGGAGCGAAACTTTTGACATCACGGACGAGATTCCAGACTTGGCGTATGCTACCAGTGACAGTGTTGAGGTTTGGACCCCCAATCTGAACGAATGCAAAAGAAGCACGAGGTTGTGTTTCTGTCGAATAAGGTCAAACGGCTGGAGAATGCCCCCGAAATCCGGAATCGTTCTCTCCTACATTAAAAATGACGAATCAAAACACTTGCAAAAAGATCATAAAACCGAAATGCTGAAGAGCTGTGAGTAA

**Class II**

**DWF-55, ENSSSA1UGG009783, SSA1-SG1 Sugar Transporter**

ATGACCAAGAAGATGGAGTCCTCAGAAAATCCAGCCGAAAGCCAGACTGAAGACCTGCTACCTAAGTCAAAAGAAATAAAATACAAAAATGCCGGCAGATCGACATTCTCACAGATCGTGGCTATGCTTGTCTTGGACTGGTTGCTCGTTGACTTCGGGCTGGAGCTGATCATACCCACCATCGTGATAGGTGCTCTTCATAAAAACCCTGCCGAAGCTCTAAACTTGACCGACGAGCAAGCATCGTGGTTTGGAAGCATCTTGTATTTCGCGCACCCGATCGGGGCTCTGATCTCAGGGTTCCTGCAGGAGCACCTGGGGCGGAAGCGGAGTCTCCTGCTGGTGAACATCCCCATGCAGGTGGCCTGGAGCACGCTGTACCTGGCCAGCTCAGTCTACCAGCTCTACTTTGTCTCGGCCACGCTTGGCCTTTGCATCGGCTTTTGCGAGGCCCCGCTCCACTCGTACATCGGCGAGATGAGCGAACCGCACGTGCGAGGCAGCCTCTCGGCCATGGGCACGGCCTCTTGCCTCATGGGCATGCTCATCATGTACCTTATCGGCTACCTCGCCCACTGGAGGACGGCCGCCTTGATCAGCAGCTTCTTTCCGGTCATTACGTTCCTCGCCATTGCCCAGATTCCAGAATCACCCACTTGGTTGGTCATGAATGGCCGCGAAAAAGAAGCCCAAAAAGCTTTGGGTTGGCTGAGAGGCTGGCTCAAGCCGGAGGAGGTTCAGGAGGAATTTCAGCGTTTGCTAGAGTACACTGACACTAAGCCAAAGTCGAATCGATTTCGCGGCTCTCAAAGAGAGAAATATGAGATGGTCCCAACCTCTGAAAACGGAGTGCCGCAACCGCTTCGAGATGATAGTGAAAGTTATTGGAGGAAAAAATTCCGGGAAATTACCAACAAAAAACTGTATTTACCACTGCGTATGGTATTTATCGTGTTCGTTTTTAGTATTGCAACTCAGCTCGCCGCTATGAGACCGTTTATGGTCGGTGTTCTCAAACAATTTGGACTTACTGTTGACAACTACTTAGTTTTGGTTTTGATCTCCTTTTTCTACTTTGTGGGCGCAATGATGAACGTGATCTTCGTGCGACGATTGGGGAAGCGACGACTCACTTTGTACTGCCAAGCAATCGCCACGCTCTCCGTTCTGCTTTTAGGCGTGTACCTCGCCTACCTCACGGGGCCAACCAAGGTCCGAGCCATTGACTGGATCCCAATTTCGCTCTTCGTCTCTGTGTTCTTCTTCAGTGGCTCCAGCATTGCTCTCATCGCTTGGCAACTTTGCGCAGAAGTTTTTCCCGTCGAAGGCCGGGGGACTGCACAAGGCCTTGTTGCAGCCTGGGCGTACTCGGTGAATTTTGTGATGTCAAAGTCCTATTTGTACTTGGAAAGGCTGGTCCAGCTCAAAGGAGTCTTTTATTTTTATGGTGCACTCTCAGTACTGGGCTTTTTCTACTATTGGCGCTATCTACCGGAGACCGAGGGCAAGTCACTTGATCAGATTGAAACTTACTTCACGGAGAACTGCGATGAGAAGGAAAAATTCACAAAACGTAAAAATAACCGTAGTTAG

**DWF-60, ENSSSA1UGG000580, SSA1-SG1 Trehalose 6-Phosphate Synthase**

ATGGGCAGTATAGGCAATGATCAACCTGCTACTGGGAAAATCATCGTGGTCGCCAATAGGTTACCGTTTGTCTTGAAACGAAATGAAGCCGGGAAACTCATTCGGATTGCCAGTGCTGGTGGTTTAGTCACAGCTGTTGCACCCGTGGTCGTACGCGGTAAAGGACTGTGGGTTGGATGGCCAGGAATTCACCTTGTAAACTTAAGTGAACCCATTCCAGAATCGGATCCAGCAGATCAAACTCCTACAGCTGGTCTTTTATCAGAAAAAGTGGTGGCTGTGCACGTAGAGCCTGCTATTTTTGATGCCTATTACAATGGTTGTTGTAATGGCACATTTTGGCCTCTATTCCACTCAATGCCGGATAGAACTATTTTCAGTGCGGATAATTGGAAAGCGTATCAGGCAGTAAATTTAGAATTTGCAGAGAAAACAGTGAATGCTTTAAGAAGGCTGATAGAGACTGAAGAAGAAAATGCTGGGACTCCTTTAGTCTGGTTACATGATTACCATCTTATGCTGGCAGCCAACACAATTCGCAATTCTGCAGATGACCTAGGATTAAAATGTAAGCTTGGATTTTTCTTACATATACCTTTTCCTCCGTGGGATATATTTCGTCTTTTCCCATGGGCAGATGAAGTGCTACAAGGAATGCTGGGATGTGATATGGTAGGCTTCCATATTGAGGACTATTGTCTGAACTTCGTTGATTGCTGCCAGCGAAGATTAGGGTGCCGTGTCGACAGGAAAGGTCTTCTTGTGGAGCATGGAGGGCGCACAGTCAGAGTTCGACCGTTACCCATTGGTATACCCTACGAAAGATTCGTCGAGTTGGCTAAGGCAGCACCACCGGTAATCACAAACAAAGATCAAAAAACCGTGCTAGGTGTCGACAGATTGGATTATACCAAAGGATTAGTCCACCGCCTCAAATCTTTTGAAGTTCTCTTAGAAAAACATCCGGAACACAGAGAAAAGGTGGTACTGATGCAGATTGCTGTTCCATCACGAACAGATGTCAAAGAATATCAGGACCTGAAGGAAGAAATGGACCAACTCGTTGGCAGAATCAATGGTCGATTCACCACACCTAATTGGTCTCCCATAAAATACATTTACGGATGTGTGTCGCAGGATGAATTGGCAGCTTTTTACAGGGAAGCTGCAGTAGCCTTGGTCACTCCTTTGAGAGACGGTATGAATTTAGTAGCAAAAGAGTTCGTTGCTTGCCAAATCAACACACCACCTGGAGTTTTGATTGTATCCCCCTTTGCTGGTGCCGGAGAAATGATGCATGAAGCTCTCATTTGTAATCCATATGAGATCAATGAAGCAGCCGAGGTCATACACAGAGCTTTAACGATGCCGGAAGATGAAAGAACTTTGCGAATGAACTCCTTGCGTAAGAGGGAAAAAGTGCATGATGTCGATTATTGGATGAGGTCCTTTTTAAAAGGAATGGGAACTCTCATCTCTGAAGATGGCGATGATGTTCTTCCAACAACAATGCAGCCTGTGACTCTGGAAGACTTCGATGAATACCTTTCCAAATACATAGGAAACACCGACAAATTGGCGCTTATGTTGGATTATGATGGTACCCTGGCCCCTATTGCTCCTCATCCAGATCTTGCCATTTTACCTCCAGAAACCAAACATGTTCTTGAAAGGCTTTCCAATATGCCTGAAGTATATATTTCAATAATCTCTGGACGGAATGTCAATAATGTTAAAGCAATGGTTGGTATCGAAGGTTTGACGTATGCTGGTAATCATGGTCTAGAAATCCTTCATCCAGATGGAAGCAGATTCGTACATCCTATGCCTGTTGAATTTGAAGACAAGGTCAGCGACCTGCTTAAGGTTCTTCAGGAGCAGGTTTGCAAGGAAGGTGCATGGGTGGAAAATAAAGGGGCTCTTTTAACTTTCCATTATCGGGAAACGCCATTAGATGTCAGACAAGAAATGGTCGCTCAAGCAAAACTTCTCATCGAACAAGCTGGTTTCAAGCCTGCCCAAGCTCATTGCGCAATCGAAGCAAAGCCGCCTGTGCAATGGAACAAAGGAAGAGCCAGTATATATATTCTACGAACAGCTTTTGGTCTAGATTGGTCGGAACGCATCCGAATCATCTATGCAGGTGATGACACCACAGATGAAGACGCCATGGAAGCATTGAAAGGAATGGCTGCTACCTTCAGAGTAACGCAATCTCAGATCATCCGCACCGTTGCAGAAAGGCGTTTGCCAAGTACAGACTCTGTTTTGACCATGTTAAAATGGGTAGAAAGACACTTCGGCAAGAGACTTCCCCGAGGAAACAGCTATGGTAGTAATCTGTGTAAGAAAGGCGATGTGATGAAAATGGAGATGAGCTTCAAACAGATGATATCTCTCCAACCAGTTAA

**DWF-66, ENSSSA1UGG010610SSA1-SG1 UTP— Glucose-1-phosphate Uridylyltransferase**

ATGACATCTCAAATCGCCGATGAATTTTTACTCGGCCACGCTGAAAAAGCTGTANNNCCAACTGACAGCCCTGATCAACCTCGATTTTATAGAAAGGTGCGAGGCCACCAACGAGTTCCCTCAGGTTCGCAAGAATTCAAAGACCTCACCAAACGTGACGCTTTGGAACAAATGAAAAACGATATCGAGCGGTTGGTTTCAACGGTGCCTGAGGCGCAGCGAGAAGCCACGAAAAAGGAGTTGGATGGGTTCTCGTCTCTATTTGAACGGTACCTCCAAGAGAAAGGTCCATCTCTCTCTTGGAATGATATTCAGAAACTTCCTGAAAATGCGGTNAGAGACTATGCCTCATTGGAAACCCCATCGACAGAAGATGCCCGGAACATGTTGGAAAAATTGGTGGTGGTCAAGTTAAATGGTGGTCTTGGAACATCCATGGGTTGCAGGGGTCCAAAATCTGTAATTCCTGTCCGGAGTGATCTTACGTTCCTAGATCTTACTGTCCAACAGATTGAATACCTAAACAAAACATACAAAGTTAATGTGCCTTTGGTTTTAATGAACTCTTTCAACACTGATGCCGATACACAGAAAATAATTCGCAAGTACAAAGGTGTTCAAGTTGAAATCTACACATTTAGTCAAAGTTGTTTCCCCAGAATAGAGAAAGAGTCGCTCATGCCTGTTCCCAAAGACTGTCATGTTGAAAAGAATTTAGAAGCGTGGTACCCTCCTGGTCATGGTGACTTTTATGAGAGCTTCAGTAACTCAGGGCTTCTCAAGAAGTTCATAAAACAGGGGCGAAAATTCTGCTTTCTCTCCAATATTGACAACTTGGGAGCCACAGTGGATCTGAATATCTTGAATATGCTGTTGAAATCAAATTCAACATCTGAATTTTTAATGGAAGTCACAGACAAAACCAGGGCTGATGTGAAGGGAGGTACCCTCATTCAGTACGACAACAAGCTGAGACTTCTGGAAATTGCACAAGTACCCAAAGAACACGTCGATGAATTTAAATCAGTGAAAACTTTCAAATTTTTCAATACAAACAATTTGTGGGTGGACCTAGAAGCCATTGAAAGAGTTGTCTCTGAGGGTGGACTAAACATGGAAATTATCATCAATCATAAGTCACTAGGAAATGGTACGAACGTAATCCAATTAGAAACAGCAGTTGGTGCCGCCATGAAAGCTTTCAATCACAGTGTTGGCTTAAATGTACCGAGGAGTCGATTCTTACCCGTCAAGAAAACATCCGACTTAATGCTAGTCATGAGTAACCTGTATGATTTAAACCATGGGTCATTGATCATGAGCCCGAAGAGAATGTTCCCGACAACTCCTCTTATCAAGCTTGGAGATGCACACTTTGCCAAAGTTAAAGATTTTCTCAGTCGTTTCGCCAATATCCCAGACTTGCTGGAGTTAGACCATTTAACCGTCTCTGGAGATGTTACATTCGGAAGAGGAGTCTCATTCAAGGGCACTGTCATAATCATCGCCAATCATGGAGACAGGATTGACATCCCAGCAGAATCACTTTTAGAAAACAAAATTATCTCCGGAAACTTGAGAATTTTGGACCACTAG

**Class III**

**DWF-61, ENSSSA1UGG013278, SSA1-SG1 Arginosuccinate lyase(ArgH)**

ATGCAAGGGTTTCAGATGGCGCTCTGGGGCGGGCGATTCTCGGAGGCGGCGGACCAGCGCTTCAAGCTCTTCAACGACTCGCTGCGCTTCGACTACCGCCTCGCGGAGCAGGACATCCTCGGCTCCATCGCTTGGTCCAAGGCTCTCGTCACCGTCAACGTCTTGACCAGCCAAGAGCAGAAGGAACTCGAGAGCGCCCTCAACGAGCTTCTGGCCGAGGTCCGGAGCCACCCCGAGCAGATACCTCAAAGCGACGCCGAGGACATACACTCCTGGGTCGAGTCTCGGCTCATCGAGAAAGTCGGGGCTTTGGGCAAGAAGCTCCACACCGGCCGCAGCCGCAATGATCAGGTGGCGACCGACCTAAAGCTCTGGTGTAAGGCACAAGTCAAGGATCTGATCCTGGCGACGCGCGGGCTACAGCAAGCGCTCATCACCACCGCCGAAGCGACCCAAGACATCCTCATGCCCGGCTACACCCACCTGCAGCGCGCTCAGCCTATCAGCTTCGCGCACTGGTGCCTGGCCTACGTGGAGATGCTGGCGCGGGATGAGACCCGTCTAAGGGACACCCTCAAGCGGCTGGACGTGAGTCCTCTGGGATGCGGGGCCCTCGCCGGCACTGCCTACCCCATCGACCGCGAGCAGCTCGCAGGCTGGCTCGGCTTCGCTTCGGCCACGAGGAATAGTCTCGACACGGTCTCGGATCGGGACCATGTCTTGGAGATTCTGTCCAACGCCGCCATCGGCATGGTCCATCTCTCGCGCTTCGCCGAGGATCTCATTTTCTTCAACACGGGCGAGGCGTCCTTCGTGGAACTGTCGGACAAGGTGACCTCGGGCTCCTCGCTGATGCCGCAGAAGAAGAACCCAGACGCCCTGGAGTTGATCCGCGGGAAGTGCGGGCGGGTTCAGGGGGCGCTGACGGGGATGCTGACAACGCTGAAGGCGCTCCCGCTGGCCTACAACAAGGACATGCAGGAAGACAAGGAAGGGCTCTTCGACGCCGTCGACACGTGGCTAGGGTGCCTCCAGATGGCAGCACTCGTCCTTGACGGCATCAAGGTGAAAAAATCTCGTTGCCGAGAGGCGGCCGAGCAAGGCTACGCCAACTCGACGGAGCTGGCCGACTACCTGGTGGGCAAAGGGCTCCCCTTCCGCGAGGCCCACCACATCGTCGGCCGGGTTGTCGTCGAGGCCATCGCCCAAGGCGTGCCCCTCGGGGCGATGTCCCTGCAGGAGCTCAAGAAATTCTCCGACGTCATCGAGGGAGACGTCTACCCCGTCCTCGAGCTCCAGTCCTGCCTCGAGAAGCGGTGCGCCAAGGGAGGCGTCGCGCCCGCCCAAGTGGCGCTGGCCATCGCCGAGGCAAAAGAACGTCTCCTCGGAAGTGAACCGTGAAAAGCGTTCTCTCTGGACTGAGAGGCTCCCTTAAAAAGGTAATTCTGCGCATTTCTGAGAACTCAGGAGTCTCTTCGACATTTGGACCGCGTTAAGGTGATTCAATGGACGCCATGTTTTTTGTCAGGACGGCATGCGATATATCGCATCAATTGGTTCCATTTTTTCCGCTACTCGTCATTTTTCTCCAAATTTGAGATCGCAATTCTGTTGTCAGGAGACCAAAGAACTCACTTACCAATTTTAACAAATAAATTCAACGTAATAAAGGCGTGGTTTTTTTTA

**DWF-63, ENSSSA1UGG005406, SSA1-SG1 Diaminopimelate decarboxylase (LysA)**

ATGGCCGGAAAAGCGCCACTTTGGAATCACTTCACGTATCGTAATAATGAACTCTACTGCGAAGATATTCCAGTCGCAAATCTTGCTGAAAAATATGGCACACCATTGTACGTATACAGCCAGGCAGCAATTCTCAAGCAGCTTCAGGCACTCCAGGAGGCATTCTCTGAGCTGAAGCCACTAGTCTGTTATTCAGTAAAGGCAAATTCCAATCTATCAATTCTTCGCTTGTTGGCGGAACATGGAAGTGGATTTGATGTGGTATCTGGTGGTGAGTTGAAGCGCGTCAGTGTCGCTGGAGGCCAGGCTAGACAAACTGTCTTTGCTGGAGTTGGCAAAACCGATGAGGAGATTGCTGCTGGCCTGACCGCTGGGGTCCTAATGTTCAACGTGGAGAGTGAGGAGGAATTGGATGCTATTGTCCGAGTAGCTCAATCCAAAAAAATCCAGGCACCTGTGGCTCTCCGAATCAATCCTGACGTTGACCCTAAAACGCATCGGTATATTTCGACAGGCCATAAGGAGTCAAAGTTCGGCATGGATATTCAACGATCATTATTGCTTGCGGAAAAATTCAAAGGCTCAGAATCTATAAAGATGATTGGCATTCATATGCATATAGGGTCGCAGATTACAACAACGACCCCTTACAGCAGTGCAGCAGCTAAGGCTGTTGATATTATTACTCGTCTGCGGGAAATAGGCCACCCAATCGACTGGTTTAACATGGGTGGCGGCTTTGGAATCAGTTACGACGGGAAGACGAGTCCATCAATTGCAGAATTTGCTGCTGCTGTTGTACCCAGCTTAAGGAAAACGGGGTGCCGTCTGGTGATGGAACCTGGTCGGGTGATTGTTGGTAATGCCGGCATTCTTATCAGTCGAGTAATTTACACAAAGGTATCAGGGGAAAAACAGTTTCTGATCCAGGACGCCGCAATGAATGATCTAATCCGTCCATCTTTATACGAGTCGTTTCATGGGATTTGGCCAGTCAAAGTCCCTGCTGGCTTTCCTTCCCCACCGGATGATTTTGAAACATCGCGTATTAATGGAACAAAATTATGCGATGTGGTTGGCCCAGTATGTGAAAGCGGTGACTTCCTGGCGCAAGATCGAGCTTTGCCACCCTTAAATCGGGGTGAGCTCCTAGCCATTTTTTCCGTTGGGGCATACGGAATGACAATGGCCTCCAACTACAATTCTCGCCCTCGCCCTGCTGAGCTGTTGGTTAATGGATCAGGCGTTGTTGTGGCTCGCCGGCGGGAGACCTTTGATGACTTAATTGCCGAGGAGCTAAATCCTGTTAGTTAA

**DWF-65,** **ENSSSA1UGT008510, SSA1-SG1 Biotin synthase (BioB)**

ATGGCCCTGAGATGGTCGCTGGAGGAGGCTTTGCGAGTGTTCAAATTACCGATGGCAGACTTGATGTATCAGGCTCAGTCTACCCACCGGGCCAACTTCAACCCTAACGAAATGCAAATCAGTACGCTTCTGAGCATCAAGACGGGCGCTTGCCCAGAAAACTGCTCGTACTGTCCCCAGTCAGCCTACTACAAAACAGAGATCAAGAAGGAGCCTCTGATGGATTTAGCGGAGGTTGTTGCCGCCGCGAAGGCAGCCAAAGAGAGTGGAAGTACCCGTTTCTGCATGGGTGCTGCTTGGCGTGGACCCACGGACAGGAATCTCCCGTTGGTCTGTGATATGGTGAAAGAGGTAAAGAAGTTGGGCTTAGAGACTTGCGTAACATTAGGGCTTCTAAAGGACCGCCACGCCGTACAACTAAAAGAAGCCGGACTGGACTTCTACAACCACAACATCGACACGTCACCGGAGTACTACAAGAAGATCATCTCGACGAGGACTTTCGAGGATCGAATCCGGACCCTGGAGCACGTGCGCAACGCAGGGATCAAGGTCTGCTGCGGCGGGATCCTAGGAATGGGTGAGAACACCGAGGACCGGATCAAGATGCTCTTGGTATTGGCGAATCTGGAGGAGCCTCCTGAATCAGTGCCGATCAATCAGCTGATTCCGATCCCTGGGACTCCTCTCGCCGACGCGGACCCTGTCGAGGGCACCGACTTCGTGCGGACCATCGCTCTCACCCGGGTCATGATGCCCAAGGCCTACATCCGACTCTCCGCTGGCCGGGAGAACATGTCCGAGGAGATGCAGACGCTCTGTTTCTTGGCAGGAGTCAACTCGATTTTCTACGGTGAGAAGCTCCTCACGGCCAAGAACTTCCGGCCGAGCAAGGATGATCAATTACTCAGTAAGCTCGGCTTCAAGAAGATGGAGATTGATGAGGCACCGGCCAACAGTCAGTGTAAAGAGGCGGCCATTGGGTGA

**DWF-62, ENSSSA1UGT027679, Chorismate Mutase (CM)**

CATAAAAGGAGCTTAAACTTAAATGTTTTATGTATTTTCCCTTTAGAAAGTAGCGCATGAAAGCAACTTATTTTCCTTTTTTTTTTTTTTTTTTTAAATTTTAATTTGTTTGTTCTGTGTGTTTCAGGTATGTAATGACAACTTTAAATTCTCTCTCCCCATTTACTCCAGCCCATTTGCTATCCTGCGACACGAGTTTGACGATGGAAACATTTTTTTCACCCTGTCAGAAGTGATGAGGGGAGCGACGAGGCAAGAGCGGACTCGCTAGTGGATGGATTCGGTTGAGCGGAAAGACAAGTCCACCGATGAGAGGAGCGGTGGATGAGCGGATGAGCCCTCGGTTCGACTGCGATGGATAGACCTTACTCGTGGGATCAATGCGGATCAGTGCTGAAGGTTGAAGCGGGGGTGGGCGGGGTCGGGGTCGCGGGGGCGGGGGAGGAAGCAAAGCCCTGGAGGACTCGGACGACGGGCCCGGGTGCTTCTGCTGCGACTCGGACAGCGACGACAGCGCCGAGCAGGTGCGGATCCGCAACACGAACAAGAAGTGCGCCCGCGGGTGCCTCAACAACATCGTCGTCTGCTCGGTCGTCATCTTGTACACCTTCTTCGGGGCCGTCCTCTTCCTCCTCATCGAGGGCGGCGGGGTCTTCACCGTCGACGACAAGATCCTCGACGAGTCCGAGGAGCCCTCCTTCCTCAACGACGAGAAGATCTCGGCCGTCAAGCGGAACCTCGGCTCCTTCAACAACCTCACGCTCCAGTGGATCGAGAACATCAACGCCGAGAACCGTGAGCGCACTGTCGAGAACATCTGGGACATCACGGTGAACTTGAACATCCTCTACCGGGAGAACTGGACCCGGTTGGCCGCCCCGGAGATCAACCGGTTCCAGCAGCAGCTCGTCGAGCGCATCACGCGGGAGGTCACCGCCTCCGGTCCCTCCGAGTCCGAGCTTCAGGTCCCGGTGCCCGTGCCGGTTCAGGTCGAACGTCACCTCGTCGACTCTTCCTATCGGATGGATCAGTATCCTCTGGAGTGGACACTCGCCAAGGCTTTCCTCTTCTCCCTCACCATCATCACGACAATAGGTGAGCATTGGACGGCCTTCACTCCCGAGCAATTTTTCAAGCCTTGGCATAAAATCGGAGCGAAAATGGGGCTCATTGAATCAGCATTAATGGCCATGGCGGTATCACTGTGCTTATTCGTCTTCGCACACTCGAAAAGCGTGGCGGTATGCCAGCCGCAAACCGCGGAAGTATGCCACCCGCAAACGGCGGAAGCGAATGCGGACGCTTCGACAGGAGCTATGATCATCCTATCCCTGGCCTCGACGCTGAACAATCGGATGGAGTACATGAAAGACGTGGCCGGGAACAAGTTCAAAGCCAACAAGCCGGTCGAGGATCTACCCCGCGAGGAGAAGGTCATGAACTCGGCCGTCGCCAACGCCTCGGCCGAGGGGCTGGATCCGAATTCGATACGGCCCTTCATCCAGGCGCAGATGGACGTGGCCAAGGACATCCAGCGCGTCTATATTGCCGGCTGGAAGGTTAGGCCAGAGAAGTGGGATCCCAGGGCGCTCGAGGAGGTCAGGGAGCTCATCTCCGCGGACGACGCCAAGATCCTGAGGCTCAGTCGACAGCAACTTACGGTTGGAGGGTTTTGCGAGGCCTATAGGGAGCAGATCTACAGGAGCCTCACAGCGCCTTACATCACTGATGCGCATAGAAAACTCCTCGTGGATACGCTGTTTTGCGTTAGGCTTAAGCAGTAG

**DWF-65, ENSSSA1UGT023765, Diaminopimelate epimerase**

CATCTTGAATTTTCACCCATCTATAAAATTACAGACATGCACGGCCTGGGCAATGACTTCATGGTCGTGGACACCGTCACCCAGAACATCTCCTTCACGACGGACCAGATCCGTCTCTTGGCCGACCGACACCTGGGCGTCGGGTTCGACCAGCTCCTGGTCGTCGAGCCCTCGTCGGAGCCCGACGTGGACTTCCACTACCGGATCTTCAACGTGGACGGAAGCGAGGTGGCCCAATGCGGGAACGGCGCCCGGTGTTTCGCCCGCTTCGTCCGGCTCCAGCGCTTGAGCCGGAAGCGCGACCTCCACGTCAGCACCGCCTCCGGGCGCCGCATGGTCCTCTCCATCACGCCGGACGACCAGGTCCGCGTCAACATGGGCGAGCCCACTCTTTACCACATGGAGAACGTGACGTTCGGTTGCGTGTCGCTAGGCAACCCGCACTGCGTGGTGCAGGTAGAGAGTGTTAAGACCGCGCCTGTGGACACGCTCGGGGCCATGCTGGAGACCCACGAGCGCTTCCCGGAGCGAACCAATGTGGGCTTTATGGAGGTCGTCGATCCCGGGCATATCCGCCTTCGCGTGTTCGAGCGTGGTGTGGGCGAGACGAGGGCCTGTGGCAGCAACGCCTGTGCTGCTGTTGTCTGCGGCGTTCGCCAGGGTGCGCTGACGTCGCCGGTGCGCGTGGATCTCCCTGGCGGCACGCTCCGTGTCGCCTGGGCGGGGCTTGGACAGCCGGTCTACTTGACCGGTCCGGCGGTTCATGTTTATGATGGCTTCGTTGATTTGGGAAAACATGA

**Class IV**

**DWF-64, ENSSSA1UGG013582, SSA1-SG1 ABC transporter**

ATGGATCACTATTTTTGCAACTCACCTCTGTGGGATTTAAATGTGACATGGAACACCAGTGATCCTAATTTCACTCCATGTCTTCAAGCAACAATTCTGTCATGGGTGCCTTGCCTATTTTTGTGGTTTTTTGCACCATTTTCAGTTCGGAGTCTAATGAAACGCCAGACATGGACAGTCCCTTGGGCTTGGCTGAATACGGCAAAATTTCTGTTGACCTTACTTTTAATATTTTTATCATTAGTTGACCTCTCTTACTCTGTTATCTCGTATTTTGACGGGAGACCATTTTATCTAGTGGATTTTATCACCCCTCTCCTAAAGACTGTCACTTTTGCTTTAGCTGCTGCTCTTCTGCTTTTGTATAAACTTCGAGGTGTCCCCTCCTCAGCCTTGTTGTTCTTGTTTTGGTTTTTGTCAGTCATCTGTGGTGCTCCACGATATAAATATGAATTGGACCGTGCTTATCGTCAGAATTTGGGCGAAGCAGTTGTTCCTTTCGTCAGCTACATGGTATATTATGCCATGGTACTTGGTTTGTTCATCCTAAATTTTTTTGCTGATCTCTTACCCATCTCACCTTCCTATTCACCGAAAGAAAAAAGTTCTCCGTACTCGCAAGCCTCCTTTCCATCATTGGTGACATTCTCATGGTTTGATCCTCTTGCATGGAAAGGTTACAGACGCCCCATTGTCGTTGAAGACTTGTGGTCTCTCGCAGAAGATGACTCTGTAAAAAAAATTGCACCTAAATTCAACAAGCATTGCAGGAAGCGATCAAATAAACATCCTCAGTCTAGAAACGTTAAGGAAGGCACAACCAGGACTGCAAATGGTATTGAGCTGAAGCCCTATTCCGACAAAGCAGCCTCCAAGGAAGTTTACCCATCCATTTGGTTCCCATTAATTAAAACTTTTTGGCCATCCGTTTTGTTTGGATTTAGTCTTAAACTTTGCAATGACATTCTCATTTTTGTGAGCCCTCATATATTGAAATCACTGATTAGCTTTATGCAAAGTGAGGAGGAAGTTTGGCATGGCTATTTCTATGCAGGAGGGCTATTTATCGCTGCTTCCTTTCAGACACTCTTCCTTGCCCATTACTTCAACCGCATGAGGATAACAGGGCTTAGGATACGAGTTGCTCTCACGTCTGCTATTTATCGTAAGGCTCTAAATATATCCAATGCATCTCGGAAAGAGTCAACAGTCGGCGAAATTGTGAATCTTATGTCTGTCGATGCGCAACGATTTATGGAACTAATGCCTCACCTATCTTTGCTTTGGTCTGCACCATTGCAGATTTCTCTGGCGGTCTTTTTCCTGTGGGAGATTTTAGGACCATCTGTCTTGGGAGGTTTAGTTGTGATGATCCTAGTAATTTTGATCAATATTCCAATTGCAAGAAAAAGTGATGCATTGACCATGAAACAAATGCGATACAAGGATGAACGTATCAAATTAATGAATGAGATTTTATCTGGAATCAAAGTGCTGAAGTTGTATGCATGGGAAACAAGCTTTGAAAAGCACATCCAGAAGATTCGAATGAAGGAGGTGAACACACTAAAACAACTCGCAATCTTAGGGTCAACAACTGCCTTCATCTGGTCTTGTGCACCACTTCTTGTAGCACTGACAACATTTGGTGTTTATTTATTCATTGACGATGAGCACTCATTAGATGCAGAAAAGACTTTTGCAACAGTGGCGTTGTTCAATATTCTCAAGTTCCCACTTGTCATGTTGCCGAACATGATCTCCAATCTTGTCCAGACAATGGTTTCTGTTAGAAGATTAAATAAGTTTATGAACTCTGATGAATTGGACCCGAATTGTGTGACACATTCTGCCTCTGAAACGGAGCCTCTGGTAATCGAAAAAGGAAATTTTGGATGGGGAACAACAGTACCTGTCCTACATGATATAGATATCAAGATTTTTGATAAGTCTCTTGTCGCAGTCGTTGGAACTGTTGGTTGTGGAAAATCTTCACTACTTGCAGCATTTCTAGGTGAAATGGATCTCATTAGCGGGAGAGTCAACACAAAGGGTTCCATTGCATATGTTACTCAGCAAGCATGGATTCAAAATGCAACTCTTCGTGAAAACATTCTATTTGGGAAGGAGTATGATTACAGAACATACAATAAGGTCATCAAAGCATGTGCTTTAAGGAGTGATTTTGCATTGTTGCCAGGTGGAGACCAAACAGAAATTGGTGAAAAAGGTATTAATTTAAGTGGAGGACAAAAACAACGAGTCAGTTTAGCTCGAGCTGTGTATAGTGATGCTGATGTATATTTTCTTGATGACCCTTTAAGTGCAGTGGATAGTCATGTGGGCAAGCACATTTTTGAAAATGTAATTGGGCCTCAGGGACTTCTTAAGAATAAAACACGGGTCCTTGTCACTCATGGATTGACCTATCTACCAGATGTGGATGAGATATATGTCTTGAAAAATGGAATGATATCAGAGAAAGGAACCTACAAACAGCTCATTGACAAAAAAGGCGCTTTTGCCGAGTTCATTCAAACCCACATGTTGGAGAGTGAAAATGATGATAAACCAGCAGAAGGAACGCAAGACATATCGCGACCGTCCTCTCCACTTGTTTATCAAAGAGTCCAAGATAGATCACGTCCCTCTTCTCCAGCCCTGTTACGTAAAAGAAGTCATAGTGCCAGTGAAAGAGGATCCACCTTGTCCTTACGGCAATTTTCGTACGAAGAAGGATCTAATTTTGTGGAGCAGGATGATGACAGGAGGAACAAGATAATTGAAGAAGAAATAGCCATGACAGGAAATATAAAATGGTCTGTCTTCTCCCATTACATGCGAGCAATGGGTTGGCTCTTTGTTTTCCTGGCATTTGCTTTTACCCTACTTTTCCAAGTTTGTGCCGTTGGGTCAAATCTTTGGCTGTCTATATGGACGTCTCGAAATAATACTCTACCAGATGGATCACAGGATCCAGAAGAAAGAAATTGGTTCCTGGAAGTCTACGCCATTCTAGGACTTTCCGCAGGTTTGTGTGCAATTGTAGCTGATCTTGCACCAAGAATTGGGTGTGTAAAAGCAAGTCGGGCTCTTCATGGTTACCTGTTACATGGTATCTTGAGAGCTCCTCTCACATTCACAGACACAACCCCTACTGGACGAATCCTTTCCCGCTTCTCGAAAGATGTAGACACTATGGATAATAAATTACCAATGGAAATATCGGACACTCTCTATTGTGTCATGGAGGTGATAGGGGTCTTGTTCATTATCACCTATGCCACACCTCCATTTTTATGGTGCATTGTGCCTTTAGTTATTCTCTACTATATTCTTCAGAGGTTTTACACGGCCACATCACGACAGTTACAAAGATTAGAGTCCATTTCGCGGTCTCCAATTTTTTCCCACTTCAGTGAAACTGTTGCAGGAGCTCAAGTTATCAGAGCATATGGAGCTGTAAACCGGTTTATTCAAGAATATGAGAAGAGAATGGATGCCAACCAAATCTGTTTCCATCCAACTGTTATATCAATCAGATGGCTGTCAGTTCGATTAGAAACTATCGGTAACTTGATAATATTTTGTGCTGCCTTTTTTGCTGTTTTAGCGCGAGGCAGCTTATCAGCTGGTCTTGTCGGCTTGTCTGTTAGCTACGCCCTTCAGGTAACTGGAGCATTGAATTGGTTTGTGCGTATGGCATCAAGTGTTGAAACAGATGTTGTTGCAATTGAGCGTGTGAAAGAGTACACTGAAATCCAAGAGGAAGCAAAATGGGATGTGCCAGAAACATTGCCTCCAGCTGAGTGGCCCCAAAAAGGAATTATTGTGTTTGAAAATTTCAAAGTTCGTTACAGGAGTGAATTGGATTTGGTTCTGAAAGGGTTGAATTTCTCTGTCAATGAAAGAGAAAAGGTCGGCATTGTGGGACGCACTGGTGCTGGAAAATCCTCATTGACCCTTGCTTTGTTCCGCATCATTGAAGCAGCAGAAGGAGCCATCTACATAGATGGCATTGATATTTCTACTCTTGGACTCCACACACTCAGATCTAGACTGACCGTCATCCCACAAGATCCAGTTCTTTTCTCTGGGAGTCTGAGGATGAATCTAGATCCGCTAGGAGTCTACTCTGATACAAAGATATGGCATGCATTAGAGTTGTCTCACTTGAAGGAGTTTGTTAAGTCCCAAGCAGCAGGTTTGCAGCATGAAATAGCTGAAGGTGGTGAAAACCTCAGTGTTGGTCAAAGGCAACTGATTTGTTTAGCCCGGGCATTGTTACGAAAAACTAAAGTACTTATTTTGGATGAAGCAACAGCAGCTGTTGATGTAGAAACAGACGATCTTATCCAAACCACCATCCGTCAGGAGTTCAGCGACTGCACAATATTGACGATTGCTCATCGGTTAAACACAATTATGGATTCCGACAGAGTTCTGGTATTGGATCACGGTCTAATGATGGAGTACGACTCTCCATTGAACCTTCTCAAGAATCCAGCGTCCCTTTTCCACTCACTCGTTAAAGATGCAGGCTTAGATGTAGAAACGATTCTCTCCACCCAAAGCATGAATGGAAGACAATTGGTGGTCTGA

**DWF67, ENSSSA1UGG011499, SSA1-SG1 UDP-glucosyltransferase2(UDP-GT2)**

ATGTTCATATTATATGAGGAACAAGATCATTGTTTGAACATGGCTTCGTCTCTGAAACTTTTGTTGGTTTCCATATTTTTCCTGGCTTCCTGTGTAAATGCATATAAGATTTTAATCTTGTATCCTACGCCTTCATACAGTCATCAAATTTCAGTTATGAACATTGCTGAAGGGCTTGTCAAGAAAGGTCATGAGGTCTTCTTCGTCAGTCCAAATTCTATATCAGGTCTGAGAGAGAATTATACATACGTGGATCTATCGTTTTCCTACAAGTATTTTAAAAAAGAGGACGATAACGAGGCTGTCAATTTAGAGCAGCAGATTTCAAGATGGGACTTCGTGAAAGTTTTTGTACCCTTCGCAGCCGTACCAGAAAAACAATTCATAAGCCAGCCTTTCGTGGAATTTCAGAGGCGGGTAGCCGCGGAGCAGATTAACTTCGACGTGGCCATCATAGAATTCGTGGGCATCCCCTACACCTGCGGTATGACGCGACTTCTGGGCAGGCTCACTAACAGCACCATCCCCTTGATCTCCATGTCCTCCATTGCGAGCGACTTCTACGCAGAGCGAAATCTAGGCAGCTTCCATCATCTGTCTTACGTGCCCACCATCTTCGACGGTTATTCGAACAAGATGTCCCTCTGGCAGAAGATCGACAACTGGTTCCTGTTTCACTGGACTGTGCTCCAGATGGAGGCGGCCCTCACAGAAAGGGCAAAGGGATTTTATAGGGATACCTACGGAGAGGCCGCTGAAGGCTTGGTGGACGGATGTTTCAGGAATTTGAGTTTGTCAGTGATATCGTCCAATGCGTTGTACTTCTATCCGAGGTTACTTGGCCCGAATGTTATTGAGATTGGACCTGTGCACTTGAAAACACCTGAAAAACTCCCAGAGAATCTGCAAAATTGGATGGACGGAGCAGAAAAAGGTGTGATTTACTTCAGTCTTGGAAGTAACATGAAAAGCAAATCTCTTCCTGCAGTTGTGCTCGATAATTTCCTGAGGTTTTTTAAACAACTTCCATCAGGCTACCGAGTCCTGTGGAAATGGGAAAAAGATGGTAATATACCAGGCCAGTCTGATAATATCTTGAGCCAAAAATGGTTACCACAACATAGCGTTCTGGCGCATCCTAAAACGAAAGTTTTCATCACTCAAGGAGGCTTGCAGAGTTTTCAAGAGGCAGTTCACTACGGAGTTCCTACGGTAGGCATACCATGGTACGGTGACCAGAATACTATCGTCGCGAAAATCCTGGATGCCCAAGTAGGCGTGCGTGTTCTCCCGAAGGAACTAACGTCATACAGTAAAATTGAATCCGGGATCAATGCTGTGCTATTTGATCGAAGATATTCGGACAATATGAAAAAACTTTCACTTATATCTCATGACTTCACCTCTAAGTCCTTGGAAAACGCTATCTTCTGGATAGAACATGTGGCAAGGCATGGTGGGGCACCTCATCTCAGGCCGTCAACAGCCGATTCTTCCTATTTTGAATTCCTATGTCTTGATATTTTATCTGTGATTCTAGTCATTACTACTCTAGTTTTGTATCTTTTGTATTTTATCTGTAAATTTACAATTGCACTGGTGTCAAATTACTTCAGTTCTAAAGAAAAAAAGTCGTGA

**DWF-68, ENSSSA1UGG008307, SSA1-SG1 UDP-glucosyltransferase 3 (UDP-GT3)**

ATGAAAACCGCAAATTTGTTAGCGATTTTAATGACATGTTTCAGCACAACTTTTTCTCATAGAGTATTAATACTTCAACCGACTCCATCATATAGTCATCAGCTCCCAGCGATGGGTTTGATAGAGGCTCTAGTGAAAAAAGGTCACGACGTTTTTTTCTTGAGCCCAGATTCAGTGCCGGGACTGCAGAAAAATTACACTCACTTCGATTTTTCATGGCTATACCAGTATTTTCACGAGAGCCTCACAGAAAAAGGTGTTCATTTACAACAACAATTTTCAAATCAACTTACCTGGATACCATTTTTGGAAGTGTGGGCAGGAACGGTGCGGCACATGTTTGAAAGTGACACTTTCAAGCAGTTTCTTAAACGCGTCCAAGATGAGAAAATATCATTCGACGTTGTGATCGTTGAGTCTTTTGTCATTCCGTACGGGGTAGGCCTGGCTCGTCTCTTGACCAAAAACAGACCTGTAATCTCCCTGGCCACCATGACTGGTGGCGAATTTTTCAACGAGGATGCCATCGGGAACATAAAACATTTATCATATTCTCCAGCAATGCTAAGCGCTTTCAATGGTAAAATGAGTCTATGGGAGCGTTTGGAAAATTGGGTCACGCATCACTACATATCGTCGAAATTGCAGAATGTTGTTGAGAATTCCGCCAAAGCCTATTTCCGAGATACTTATGGCCCAGATGGTGAGGCTTTCGTCGAAGGGTGCTGGGGAAACATTAGTCTCGCCATGATAACCTCAAACGCTCTGTATTACTATCCAAGGCCTATCACTCCCAATGTTATCGAGGTTGGGCCGCTTCATATAAAAAATTCGACAAAATTACCAAAGACATTGCAAGATTGGTTAGACGGGGCAGAACGAGGAGTAATTTATTTCAGTCTCGGCAGTAACATGAGAAGTGCAAACCTACCCCGAGAGGCATTGTCGAACTTCTTGAGGGTTTTCAGAGAGCTGCCGCAAGGTTACCGAGTGCTGTGGAAATGGGAGGACGATGCCGAAATTGCAAAAAATGAAGAGGGAGTCGGGAAATCAAAAGTAGCTAATATTCTTACTCAAAAATGGATACCACAGCAAGAAGTTTTAGCGCATCCAAAGGTAAAATTGTTCATCACACAAGGAGGATGTCAAAGCTTCCAAGAAACAGTTCACTACGGAGTACCATTGGTAGGGGTCCCTTGGTTTGGAGACCAAGAGGTGAACGTTGCAAAGATGGTGGACGCAGGAATTGGAGTTCGGCTCCGACCTCAGGAGTTGGATTCTTTTGCGAAAGTCAAACAAGCTATTAAAGCTGTATTGTTCGATAAAAGTTATGCTGAAAATATGAAGAAATTATCTGATTTATCTCATGACTTCACAAGACGAGCTCCAGATGAAGCTGTTTTTTGGGTGGAGCATGTGGCAAAACATGGTGGAGCTGCTCATTTGAGACCTTACATAGCTGATATTTCTTACTTCACATACTTCTGTCTCGATATCATATCTATTATCTTAGGAAGCAGCATAGCAATATTGTACGTTCTGTGGACTATATTTAGATAT

TTAATTTCATTAATGCCAACACTGTCAAGTAAAAAAATTAAAAACTTGTGA
