## Supplementary material for "Creating resistance to the whitefly *Bemisia tabaci* in cassava through RNAi-mediated targeting of multiple insect metabolic processes": Table S2

| **Strategy** | **Function** | **Target gene(s)** | **Accession number** | **Construct number** | **# independent transgenic events produced in cv. NASE 13** |
| --- | --- | --- | --- | --- | --- |
| **I.** Osmoregulation | Maintain osmotic balance in gut | Aquaporin (AQP) | ENSSSA1UGG010352 | DWF 11 | 14 |
|  |  | Glycoside hydrolase family 13_1 + Aquaporin | ENSSSA1UGG011937& ENSSSA1UGG010352 | DWF 15 | 9 |
|  |  | Glycoside hydrolase family 13_5 | ENSSSA1UGG000414 | DWF 57 | 9 |
|  |  | Glycoside hydrolase family 13_12 | ENSSSA1UGG002701 | DWF 59 | 5 |
| **II.** Carbohydrate Homeostasis | Energy balance, sugar transport & storage | Sugar Transporter | ENSSSA1UGG009783 | DWF 55 | 13 |
|  |  | Trehalose 6 phosphate | ENSSSA1UGG000580) | DWF 60 | 10 |
|  |  | UTP--glucose-1-phosphate uridylyltransferase | ENSSSA1UGG010610 | DWF 66 | 15 |
| **III.** Symbiosis (Nutrientbiosynthesis) | Amino acid & biotin synthesis | Arginosuccinate lyase | ENSSSA1UGG013278 | DWF 61 | 11 |
|  |  | Chorismate mutase | ENSSSA1UGT027679 | DWF 62 | 7 |
|  |  | Diaminopimelate decarboxylase | ENSSSA1UGG005406 | DWF 63 | 10 |
|  |  | Diaminopimelate epimerase | ENSSSA1UGT023765 | DWF 64 | 2 |
|  |  | Biotin synthase | ENSSSA1UGT008510 | DWF 65 | 6 |
| **IV.** Detoxification | Detoxify plant toxins  (Phase II & III detox pathways) | ABC Transporter_26 | ENSSSA1UGG013582 | DWF 56 | 7 |
|  |  | UDP glucuronosyltransferase _3 | ENSSSA1UGG011499 | DWF 67 | 11 |
|  |  | UDP-glucuronosyltransferase _2 | ENSSSA1UGG008307 | DWF 68 | 11 |
| Control | Control | GFP |  | DWF 17 | 4 |
