## Supplementary material for "Creating resistance to the whitefly *Bemisia tabaci* in cassava through RNAi-mediated targeting of multiple insect metabolic processes": Table S3

**Table S3-** **qRT-PCR primers used in this study**

| **Gene construct and target gene** | **Primer name** | **Sequence** | **Tissue** |
| --- | --- | --- | --- |
| **DWF 55**, ENSSSA1UGG009783, Sugar Transporter | Forward | CCATTGCCCAGATTCCAGA | Adults/Nymphs |
|  | Reverse | CCTCTCAGCCAACCCAAAG | Adults/Nymphs |
| **DWF 56**, ENSSSA1UGG013582, ABC Transporter_26 | Forward | TTGTGGAGCAGGATGATGAC | Adults/Nymphs |
|  | Reverse | CAAAGAGCCAACCCATTGC | Adults/Nymphs |
| **DWF 57**, ENSSSA1UGG000414, Glycoside hydrolase family 13_5 | Forward | TGCTGGAGGAAATGGTGTTC | Adults/Nymphs |
|  | Reverse | ACAGTGAGGCTGAATCTTGG | Adults/Nymphs |
| **DWF 61**, ENSSSA1UGG013278, Arginosuccinate lyase | Forward | TTTCTTCAACACGGGCGAG | Adults/Nymphs |
|  | Reverse | GTCTGGGTTCTTCTTCTGCG | Adults/Nymphs |
| **DWF 62**, ENSSSA1UGT027679, Chorismate mutase | Forward | AATAGGTGAGCATTGGACGG | Adults/Nymphs |
|  | Reverse | GCGAAGACGAATAAGCACAG | Adults/Nymphs |
| **DWF 65**, ENSSSA1UGT008510, Biotin synthase | Forward | GCCAACTTCAACCCTAACGA | Adults/Nymphs |
|  | Reverse | AACAACCTCCGCTAAATCCA | Adults/Nymphs |
| **DWF 66**, ENSSSA1UGG010610, UTP--glucose-1-phosphate uridylyltransferase | Forward | GACTTGCTGGAGTTAGACCA | Adults/Nymphs |
|  | Reverse | GTGATTCTGCTGGGATGTCA | Adults/Nymphs |
| **DWF 67**, ENSSSA1UGG011499, UDP-glucuronosyltransferase_3 | Forward | GAGTCCTGTGGAAATGGGAA | Adults/Nymphs |
|  | Reverse | AGCCTCCTTGAGTGATGAAA | Adults/Nymphs |
| Control (RPLA) | Forward | CATTCCACTACAGAGCTCCA | Adults/Nymphs |
|  | Reverse | TTTCAGGTTTCGGATGGCTT | Adults/Nymphs |
| **DWF 11**, ENSSSA1UGG010352, Aquaporin | Forward | TAGTTGCTGAGTTCGTTG | Plant leaf |
|  | Reverse | TTACAGCCGGATTGATGT | Plant leaf |
| **DWF 57**, ENSSSA1UGG000414, Glycoside hydrolase family 13_5 | Forward | GTGATTGTTGACTTTGTGCCGA | Plant leaf |
|  | Reverse | TCCGTTTACCGTCCTTCCAA | Plant leaf |
| **DWF 59**, ENSSSA1UGG002701, Glycoside hydrolase family 13_12 | Forward | GGGTGCCTCCATACAATGAT | Plant leaf |
|  | Reverse | GCTGAAGACTGATGCCCAAT | Plant leaf |
| **DWF 55**, ENSSSA1UGG009783, Sugar Transporter | Forward | TTGGACTGGTTGCTCGTTG | Plant leaf |
|  | Reverse | GCTCGTCGGTCAAGTTTAGA | Plant leaf |
| **DWF 60**, ENSSSA1UGG000580, Trehalose 6 phosphate | Forward | ATGTGTGTCGCAGGATGAAT | Plant leaf |
|  | Reverse | CAGGTGGTGTGTTGATTTGG | Plant leaf |
| **DWF 66**, ENSSSA1UGG010610, UTP--glucose-1-phosphate uridylyltransferase | Forward | ATGTGTGTCGCAGGATGAAT | Plant leaf |
|  | Reverse | CAGGTGGTGTGTTGATTTGG | Plant leaf |
| **DWF 61**, ENSSSA1UGG013278, Arginosuccinate lyase | Forward | GAGACCGTGTCGAGACTATT | Plant leaf |
|  | Reverse | GTCTAAGGGACACCCTCAAG | Plant leaf |
| **DWF 62**, ENSSSA1UGT027679, Chorismate mutase | Forward | ACTTCTCTGGCCTAACCTTC | Plant leaf |
|  | Reverse | ATTCGATACGGCCCTTCAT | Plant leaf |
| **DWF 63**, ENSSSA1UGG005406, Diaminopimelate decarboxylase | Forward | CATCCAATTCCTCCTCACTCT | Plant leaf |
|  | Reverse | CAGGCTAGACAAACTGTCTTTG | Plant leaf |
| **DWF 64**, ENSSSA1UGT023765, Diaminopimelate epimerase | Forward | GGATCGACGACCTCCATAAA | Plant leaf |
|  | Reverse | GGTGCAGGTAGAGAGTGTTA | Plant leaf |
| **DWF 65**, ENSSSA1UGT008510, Biotin synthase | Forward | GTTGACTCCTGCCAAGAAAC | Plant leaf |
|  | Reverse | ATGCCCAAGGCCTACAT | Plant leaf |
| **DWF 56**, ENSSSA1UGG013582, ABC Transporter_26 | Forward | CAGACACTCTTCCTTGCTCAT | Plant leaf |
|  | Reverse | TGTTGACTCTTTCCGAGATGC | Plant leaf |
| **DWF 67**, ENSSSA1UGG011499, UDP-glucuronosyltransferase_3 | Forward | GTGGCCAGGGAGATTACA | Plant leaf |
|  | Reverse | CGTTGTGATCGTTGAGTCTTT | Plant leaf |
| **DWF 68**, ENSSSA1UGG008307, UDP-glucuronosyltransferase_2 | Forward | GCCCTTCAGCAATGTTCATAA | Plant leaf |
|  | Reverse | TCCTGGCTTCCTGTGTAAAT | Plant leaf |
| **DWF 17**, GFP | Forward | ACCCTCTCCACTGACAGAA | Plant leaf |
|  | Reverse | ACGATGGGTAAAGGAGAAGAAC | Plant leaf |
