## Supplementary material for "Creating resistance to the whitefly *Bemisia tabaci* in cassava through RNAi-mediated targeting of multiple insect metabolic processes": Figure S1

### Slide 1
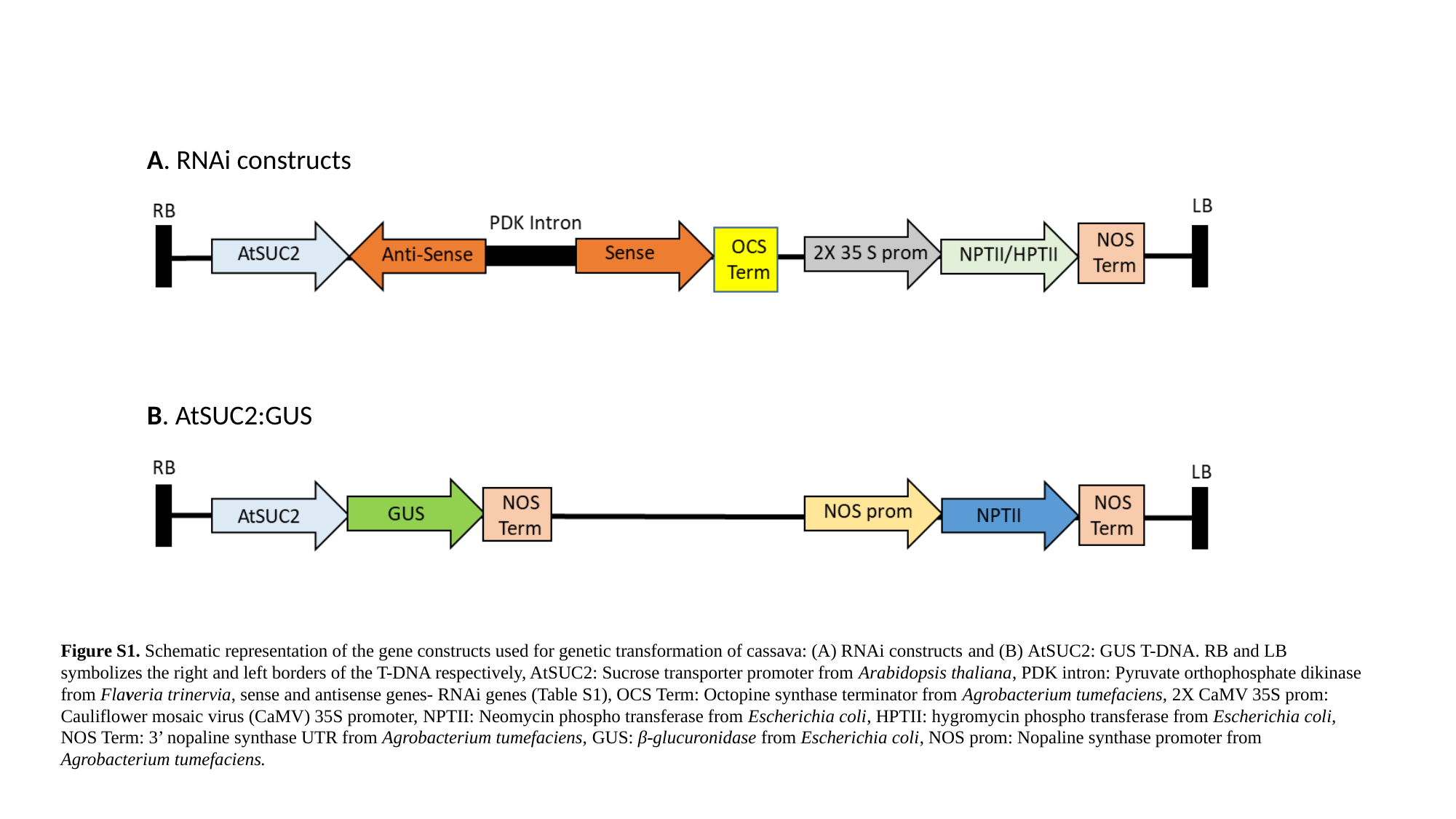

A. RNAi constructs
B. AtSUC2:GUS
Figure S1. Schematic representation of the gene constructs used for genetic transformation of cassava: (A) RNAi constructs and (B) AtSUC2: GUS T-DNA. RB and LB symbolizes the right and left borders of the T-DNA respectively, AtSUC2: Sucrose transporter promoter from Arabidopsis thaliana, PDK intron: Pyruvate orthophosphate dikinase from Flaveria trinervia, sense and antisense genes- RNAi genes (Table S1), OCS Term: Octopine synthase terminator from Agrobacterium tumefaciens, 2X CaMV 35S prom: Cauliflower mosaic virus (CaMV) 35S promoter, NPTII: Neomycin phospho transferase from Escherichia coli, HPTII: hygromycin phospho transferase from Escherichia coli, NOS Term: 3’ nopaline synthase UTR from Agrobacterium tumefaciens, GUS: β-glucuronidase from Escherichia coli, NOS prom: Nopaline synthase promoter from Agrobacterium tumefaciens.
