## Supplementary material for "Creating resistance to the whitefly *Bemisia tabaci* in cassava through RNAi-mediated targeting of multiple insect metabolic processes": Figure S2

### Slide 1
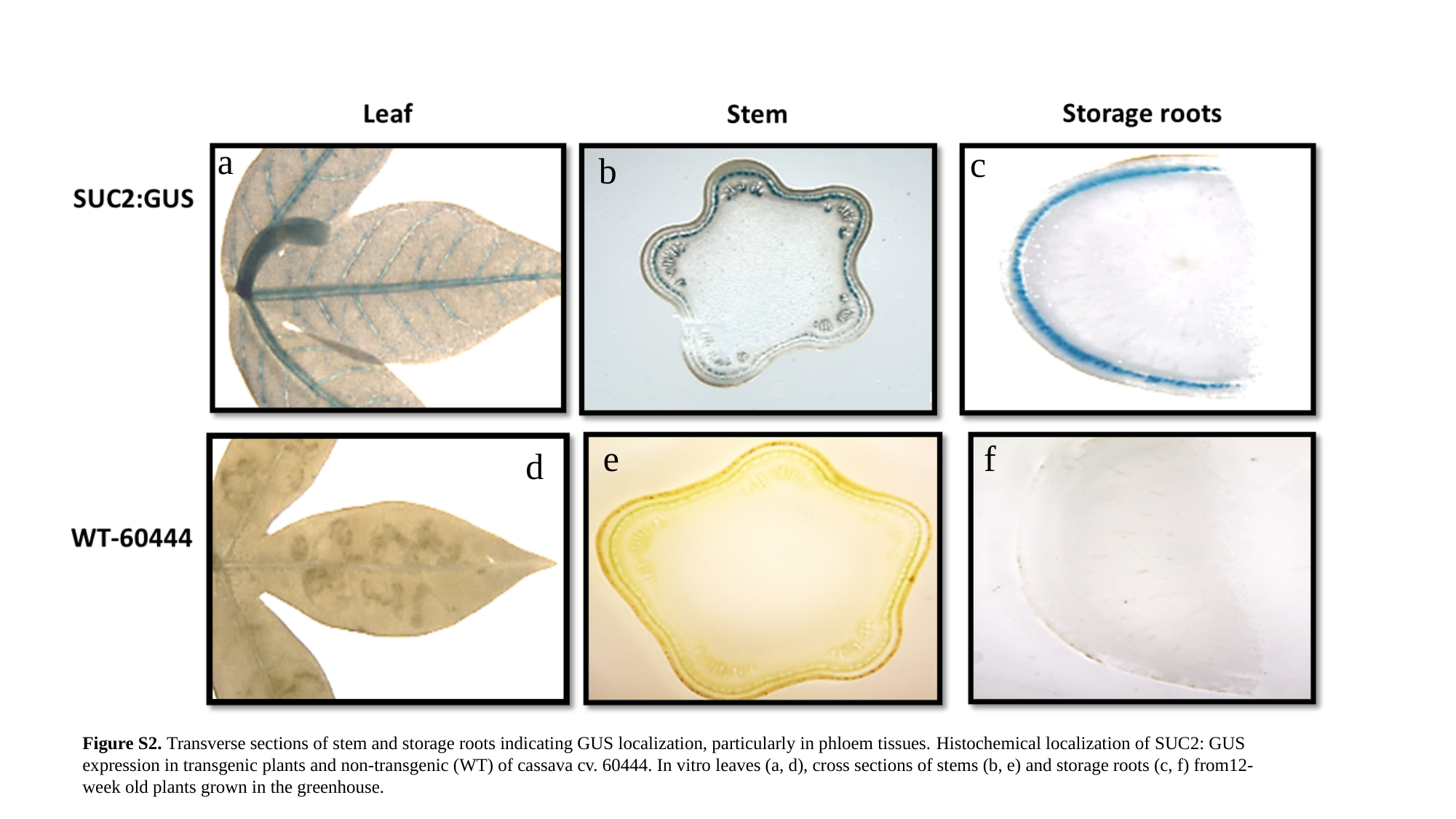

a
c
b
f
e
d
Figure S2. Transverse sections of stem and storage roots indicating GUS localization, particularly in phloem tissues. Histochemical localization of SUC2: GUS expression in transgenic plants and non-transgenic (WT) of cassava cv. 60444. In vitro leaves (a, d), cross sections of stems (b, e) and storage roots (c, f) from12-week old plants grown in the greenhouse.
